# Sequence determinants of pathogenicity in glucose-6-phosphatase linked to glycogen storage disease type 1a

**DOI:** 10.64898/2026.07.27.741017

**Authors:** Richard A. Stein, Emily M. Hawes, Chanel M. Norphlet, Margaret H. Rakonick, Shelby A. Harris, Trisha Sivam, Audrey M. Lucerne, Valentina R. Da Silva, Richard M. O’Brien, Derek P. Claxton

## Abstract

Glycogen storage disease type 1a (GSD1a) is an autosomal recessive Mendelian disorder that can be caused by missense variants in glucose-6-phosphatase catalytic subunit 1 (G6PC1). Although hundreds of missense variants have been identified, the vast majority are of unknown clinical significance, and the molecular mechanism(s) of *bona fide* pathogenic variants are ill-defined. We combine bioinformatic data and clinical associations with the protein language model AlphaMissense to guide mechanistic exploration of 78 missense variants at 55 residue positions using robust biochemical and biophysical assays to distill general principles of enzyme dysfunction. Correlation analysis established a strong linear relationship between folded G6PC1 abundance and catalytic capacity for most variants. Pathogenic variants within this paradigm were linked to compromised stability and activation of the unfolded protein response. However, outliers characterized by relatively high abundance, yet low activity clustered to a network of sidechains adjacent to the active site that allosterically modulate catalysis. Contextualized by recent high-resolution structures and AlphaFold modeling, our holistic analysis of G6PC1 *in vitro* metrics facilitates clinical (re)classification of variants according to explicit molecular phenotypes and identifies therapeutic design directions. Moreover, our approach illustrates a blueprint for variant characterization that integrates computational prediction with experimental validation to discover disease etiology.

## Introduction

The rapid expansion of whole genome and exome sequencing databases and investment in biobank repositories is uncovering the scale of human genetic diversity and facilitating discovery of genotypic links to disease (1–5). The substitution of single amino acids, or missense variants, arising through non-synonymous single nucleotide variation (SNVs) is a major contributor to genetic diversity within protein coding regions and to Mendelian (monogenic) disease (3,6). Although identification of genotypes is essential to the promise of precision medicine (7), only a small fraction of the ∼16 million missense variants observed across more than 800,000 exomes or genomes have been classified as tolerated (“benign”) or disease-causing (“pathogenic”) (8). Consequently, most SNVs are labeled as a variant of unknown significance (VUS). While standards are in place for classifying variants (9), the elucidation of causal molecular mechanisms remains a fundamental challenge in the interpretation of disease etiology and implementation of treatment strategies. As amply illustrated by channelopathies (10,11), missense variants may not only perturb function but also alter other fundamental properties such as stability or trafficking. However, the arduous task of interpreting the biochemical significance of non-synonymous SNVs is complicated often by the lack of reproducible, standardized assays that clearly define the functional landscape of affected genes.

Strategies to bridge the genotype-phenotype gap have largely been focused on developing either reporter assays combined with deep mutational scans of target proteins (12,13) or *in silico* approaches informed by sequence space and/or other tangible features (14,15). While site-specific functional constraints derived from high-throughput saturation mutagenesis have the power to uncover relevant protein properties (16,17), cell-based variant effect assays demand a significant time and resource investment that requires an assay design that fits a chosen detection scheme. In addition, not all targets are amenable to this approach. Interpreting the derived “fitness” score is not trivial as the link between target molecular properties, cell survival, and pathogenesis is not always direct and may not reflect all facets of variant perturbations (18). Moreover, the mechanistic consequences of variant perturbations are likely to vary across the proteome. For example, a recent study of 522 independently folded domains comprised of ∼100 amino acids found that while protein stability is a major contributor to pathogenesis, the relative proportion of pathogenic destabilizing variants differs across protein families (19). This suggests that while broad principles of perturbed molecular processes can be ascertained from sampling the proteome, specific gene-level features that dictate functional landscapes, and ultimately clinical decisions, are limited. From a practical perspective, systematic evaluation of thousands of full-length genes linked to disease is an immense challenge for deep mutational scans.

Though easily scalable, the predictive power of many *in silico* variant effect tools that leverage sequence and/or structural information is hampered by low to moderate correlations with experimental metrics and highly variable performance as binary classifiers (20–23). These tools vary in algorithmic complexity and either infer a phenotypic or clinical outcome (i.e. SIFT (24,25), EVE (26)) or change in protein stability (i.e. Rosetta DDG (27), FoldX (28)) caused by mutations in the primary sequence. Notably, machine learning tools trained on evolutionary sequence space with augmentation by other features has been shown to improve prediction accuracy against benchmarks (29–31), potentially distinguish roles of conserved residues in proteins (32), and even infer catalytic parameters (33). AlphaMissense is a protein language model built on the revolutionary AlphaFold2 structure prediction pipeline that combines evolutionary sequence constraints, structural context, and allele frequencies for inferring missense variant pathogenic probability on a 0 (likely benign) to 1 (likely pathogenic) scale (34). While correlations and predictability between AlphaMissense scores and experiment are dependent on assay design and robustness (34), a recent benchmark study has shown that AlphaMissense is among the best performing variant effect predictors out of 65 tested using three distinct datasets (35). Two recent studies showed that incorporation of AlphaMissense scores improved interpretation of missense variants in p53 (36) and BRCA1 (37). Despite increasing sophistication, AlphaMissense and similar predictors are not accurate enough as a lone diagnostic for genetic disease (9,38). However, such tools can be used to support preliminary phenotype class assignments (i.e., likelihood of disease) in conjunction with other data sources. Nevertheless, the mechanistic link between prediction and phenotype remains unrealized. Recently, Chang-Gonzalez et al demonstrated that integration of AlphaMissense scores into a gene-specific predictor trained on a trove of high-quality biophysical data improved variant classifications for the K+ channel KCNQ1 implicated in congenital long QT syndrome (39). In the absence of suitable experimental datasets that outline functional landscapes for disease-linked targets, this suggests that AlphaMissense could be used as a coarse map for direct mechanistic investigations.

Here, we combine variant characteristics identified in genetic databases with variant effect predictions from AlphaMissense to guide mechanistic inquiry of missense substitutions in glucose-6-phosphatase catalytic subunit 1 (G6PC1), an ER-resident membrane protein that catalyzes the hydrolysis of glucose-6-phosphate (G6P) to glucose and inorganic phosphate (Pi) (40,41). As the terminal enzyme in glucose production from glycogen or pyruvate, disruptive mutations in G6PC1 cause the autosomal recessive glycogen storage disease type 1a (GSD1a) that leads to severe, and even lethal, consequences and for which there is no curative therapy (42–44). Though rare (0.001% population frequency), hundreds of variants have been documented in genetic databases (7,8,45,46), yet clinical and mechanistic implications remain largely unknown. Foundational studies suggested disparity between enzyme expression patterns and catalytic activity (47). Our previous study of GSD1a-linked variants in the G6PC1 active site highlighted multiple, non-mutually exclusive pathways for enzyme dysfunction, such as active site collapse and reduction of G6PC1 stability (48). Informed by AlphaMissense prediction patterns of variants identified in ClinVar, gnomAD, and the Human Gene Mutation Database (HGMD), we investigated a panel of G6PC1 variants that sample more broadly a range of sequence and structural features to uncover fundamental molecular properties contributing to GSD1a etiology. Compiled from biophysical and functional assays that fingerprint G6PC1 variants *in vitro* and *in situ*, our study delineates the relationship between enzyme destabilization, abundance and catalytic capacity. Interpreted within the context of enzyme structure, variant-induced perturbations of hydrolysis kinetics invoke an allosteric model mediated by a conserved network of sidechains that modulate access to the active site. Collectively, our analysis captures deterministic features of sequence space that not only impart mechanistic inferences but also support the clinical (re)classification of G6PC1 variants.

## Results

*Pathogenicity prediction patterns in G6PC1*. We have tabulated 549 non-synonymous SNVs giving rise to missense variants at 292 amino acid positions in canonical G6PC1 (Table S1), which were assembled primarily from gnomAD_v4.1, ClinVar, and the HGMD. Features of these variants, namely allele frequency and sequence conservation, were used to parse AlphaMissense predictions. Variant data from the Regeneron Genetics Center Million Exome (RGC) project (7) reported similar allele frequency data as gnomAD_v4.1. Mapping population-level allele frequencies from gnomAD_v4.1 and the RGC study for 493 of these SNVs onto the G6PC1 model highlights that a considerable number of amino acid positions display higher substitution frequencies than others (Fig 1A), although the phenotypic and biochemical consequences are not universally known. Within ClinVar, the overwhelming majority (87%) of all variants are of uncertain clinical significance or have received conflicting interpretations of pathogenicity. Only four variants have been assigned a “benign” label, whereas 68 have been identified as “pathogenic” and causal for GSD1a. None of the variants have been extensively reviewed by an expert panel.

**Figure 1.**
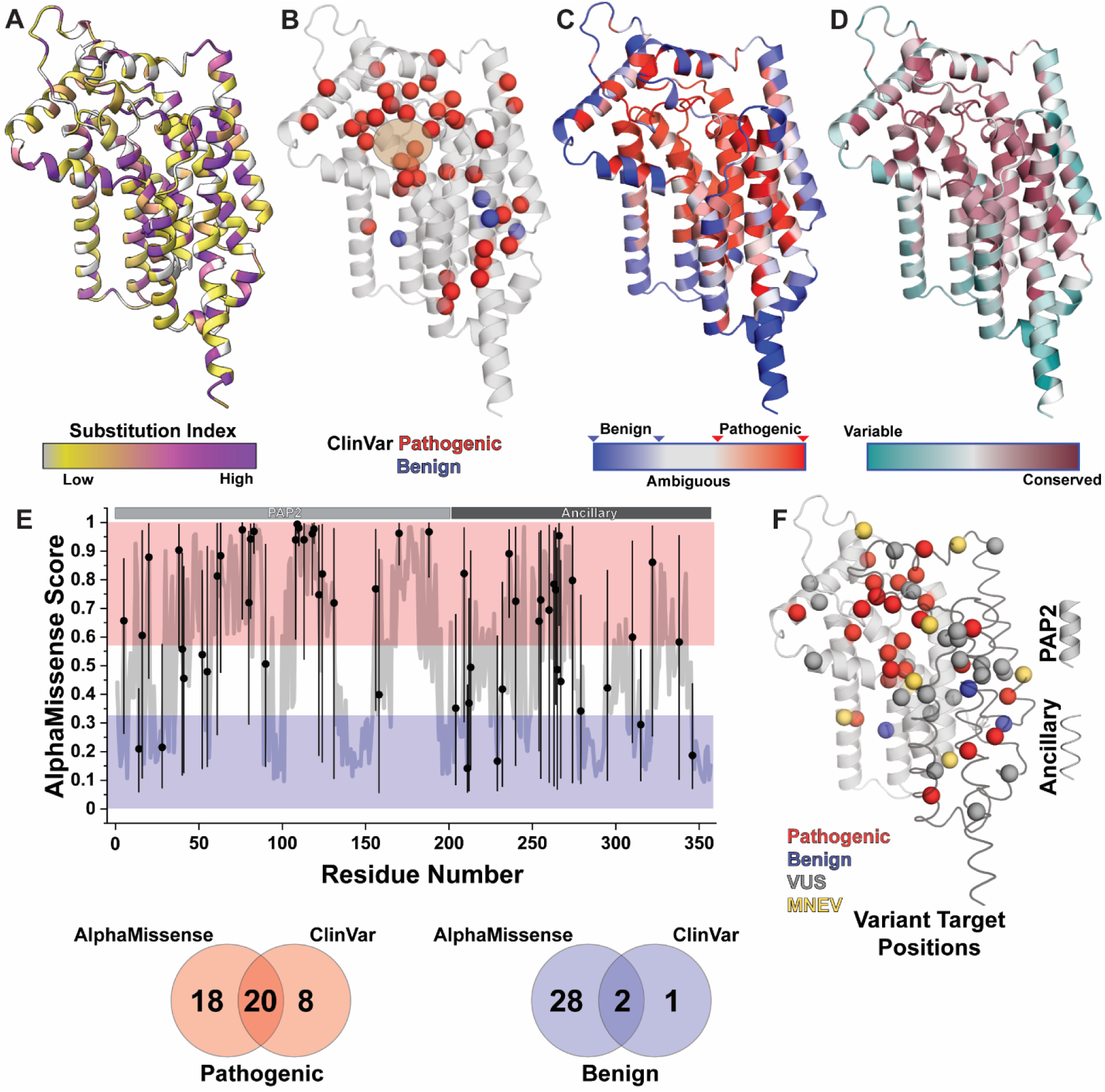
G6PC1 variant selection strategy. **A.** The substitution index represents the composite alternate allele frequency compiled from gnomAD_v4.1 and Regeneron Genetics Center (Methods), which is mapped onto a structure of G6PC1. A variant position is colored grey if allele frequency is not known. The color scale captures the range of known allele frequencies over three orders of magnitude (6.08 x 10^-7^−7.63 x10^-4^). **B.** Relative locations of GSD1a-linked variants and benign variants listed in ClinVar. The active site is highlighted. **C.** The tertiary distribution of mean pathogenic probability scores from AlphaMissense for each position shows general correspondence to the sequence evolution rate calculated by ConSurf (**D**). **E.** The average AlphaMissense score as a function of residue number is shown as a thick grey line. Positions for variants chosen for analysis are shown as black circles at the mean and the range of scores predicted for native amino acid substitution as a thin black line. The colored regions illustrate the suggested cutoffs for likely benign (blue) and likely pathogenic (red) interpretations. The similarity between predicted (AlphaMissense) and labeled (ClinVar) variant classifications are shown in the Venn diagram below. **F.** The relative positioning of variants explored in this report is shown on a structure of G6PC1 with identified PAP2 and ancillary regions. The variants are color coded by ClinVar classification, though some positions are represented in other classifications as well (Table S2). For simplicity, ClinVar labels “pathogenic” or “likely pathogenic” are represented as “Pathogenic”; “benign” or “likely benign” as “Benign”; “uncertain significance” or “conflicting interpretations” as “VUS”. Variants generated by exchanging multiple nucleobases in a codon are referred to as “MNEV”.

While the limited distribution of labeled “benign” variants displays no clear structural pattern, most “pathogenic” variants are clustered within or near the G6PC1 active site (Fig 1B). To identify amino acid substitutions that are likely to have biological consequences, we mapped the predicted average AlphaMissense score for each position onto the G6PC1 structure and color coded the scores according to the prescribed thresholds for “likely benign” and “likely pathogenic (34)”. As expected, substitution of residues that mediate packing interactions within the protein core, including the active site, are predicted to be disruptive whereas those along the surface projecting into either the aqueous or lipid phases are expected to be largely tolerated (Fig 1C). Zones of uncertainty, classified by AlphaMissense as “ambiguous”, are scattered throughout the sequence. The global scoring pattern reflected closely the sequence evolution rate calculated by ConSurf (49,50) (Fig 1D), which is expected since multiple sequence alignment is included in the AlphaMissense model architecture.

However, decomposition of residue-specific features emphasized that “likely benign” and “likely pathogenic” regions were not equivalent in predictive confidence. We noticed that the spread of AlphaMissense scores per residue informed patterns of AlphaMissense classification (Fig. S1a,b) and sequence conservation (Fig. S1c). Those sequence positions classified as “likely benign” or “likely pathogenic” on average demonstrated a broadly distributed range, a simple indicator of variability in the scoring of 19 amino acid substitutions. In contrast, the “ambiguous” class displayed a high scoring range that was more tightly distributed (Fig. S1b). Furthermore, this class included intermediate levels of sequence conservation relative to “likely benign” (less conserved) and “likely pathogenic” (more conserved) classes (Fig. S1c). A direct comparison of sequence conservation and score range illustrated that AlphaMissense classes occupy diffuse nodes in the feature space (Fig. S1d). These features of the “ambiguous” class reduce predictive confidence for any given substitution. As discussed below, this patterning helped to rationalize erroneous pathogenicity predictions. In addition, mapping of score range onto the G6PC1 structure highlighted that the highest predictive confidence (lowest scoring variability) was found in the active site (“likely pathogenic” class) and accessible loops (“likely benign class”). In contrast, the lowest confidence regions for these two classes were found within the transmembrane helices and secondary structures exposed to the ER-lumen (Fig. S1e). This suggests that the AlphaMissense score spread could operate as a heuristic for functionally important sites in target proteins.

At the level of individual SNVs, AlphaMissense concurred with 56 of the 68 labeled “pathogenic” or “likely pathogenic” variants in ClinVar. Thus, the granularity within the AlphaMissense algorithm and alignment with bioinformatic and clinical data instilled confidence that pathogenicity scores can be a useful metric to guide selection of variants for further study. Nevertheless, we found that conflicts with clinical assignments could be linked to sequence conservation patterns and relatively elevated allele frequencies. While most ClinVar-labeled pathogenic SNVs are very rare (AAF < 1E-5) with high pathogenic probability, we found that 10 scored into the “ambiguous” class (Fig. S1f). On average, this uncertainty was associated with intermediate sequence conservation and a broad range of AlphaMissense scores for the native amino acid in question (Fig. S1b,c). We also observed that several pathogenic variants linked to GSD1a displayed allele frequencies > 1E-5, the benign cutoff for the AlphaMissense model. In fact, two putatively disease-linked variants, R295C/H (44,51), with elevated allele frequencies were scored as “likely benign” by AlphaMissense. The high allele frequency of disease-linked variants is a drawback of methods that rely on allele frequency to infer pathogenicity (9,52), especially since variants can confer an environmental advantage and thus be more representative in specific human populations as exemplified by hemoglobinopathies (53,54). Therefore, we emphasize that caution must be taken when interpreting monogenic disease probability based on variant prevalence or *in silico* predictions.

*Selection of G6PC1 variants for investigation.* For reference, G6PC1 can be divided into two structural regions. The PAP2 fold consists of the first five transmembrane helices and includes the signature tripartite sequence motif (KX6RP---PSGH---SRX5HX3D) that defines the PAP2 superfamily (55–57). The last four transmembrane helices, termed ancillary helices, are unique among PAP2 superfamily membrane proteins based on a Foldseek (58) search. This global architecture is conserved in two other G6PC isoforms, G6PC2 and G6PC3, that display different tissue expression patterns (59). For this study, we selected 78 variants at 55 residue positions (highlighted as black dots) that broadly sample unique packing environments and pathogenic probabilities within both the PAP2 fold and ancillary helices (Fig 1E). Concordance of AlphaMissense scores with ClinVar labels for expected pathogenic and benign variants among the selection are indicated by the union of a Venn diagram in Fig 1E. The three-dimensional relationship of these targeted variants by amino acid position is shown in Fig 1F, color coded by their clinical classification. Detailed in Table S2, our selection included multiple SNVs at some positions, reflecting both the range of AlphaMissense scores (solid black bars, Fig 1E) and the substitution index (Fig 1A). Our targets also incorporated multi-nucleobase exchange variants (MNEVs) leading to amino acid substitutions that are unlikely to arise in the human population instantaneously (i.e., R40H: AGG→CAT/C). In some cases, these more extreme substitutions are observed in other metazoan species or other G6PC isoforms (Table S2). The inclusion of MNEVs stratifies the dataset and is intended to explore the dependence of G6PC1 abundance, stability, and activity on sidechain physiochemical properties.

*Correlation of biochemical metrics with variant effect predictors*. We screened the selected variants for enzyme abundance and hydrolysis activity following plasmid transient transfection of adherent HEK293SG cells. In these assays, G6PC1 was expressed as a fusion protein with EGFP on the C-terminus. Folded protein abundance was quantified by the area under the elution curve acquired from EGFP fluorescence detection size exclusion chromatography following G6PC1 extraction with lauryl maltose neopentyl glycol (LMNG) detergent. G6PC1 enzyme activity in LMNG micelles was measured by the amount of inorganic phosphate (Pi) released following hydrolysis of a saturating concentration of G6P. We have shown previously that these *in vitro* assays using solubilized G6PC1 distinguish active site variant properties and accurately reflect purified enzyme kinetics (48,60). Histograms of variant-dependent hydrolysis activity and abundance are shown in Fig. S2.

The predictability of AlphaMissense scores was assessed by correlation plots against the biochemical metrics. Variant AlphaMissense scores demonstrated a strong negative correlation with activity, which was captured statistically by Spearman’s rank correlation coefficient (Fig 2A). Although GSD1a-linked variants displayed a broad range of AlphaMissense scores, these were limited to ≤ 40% of WT activity. An opposite pattern was observed for the three benign-labeled variants screened. Approximately half of the variants of uncertain significance scored in the “likely benign” category and retained > 50% of WT activity, which was similar to the activity of known benign-labeled variants from ClinVar. Interestingly, comparable correlations were obtained for evolutionary model of variant effect (EVE) (26) and cross-protein transfer 1 (CPT-1) (61) predictions, two other variant effect predictor models of distinct complexity. However, both models were not as accurate as AlphaMissense in separating “benign” from “pathogenic” variant ClinVar labels, and CPT-1 scoring was highly compressed which precluded interpretation (Fig. S3, Table S1).

**Figure 2.**
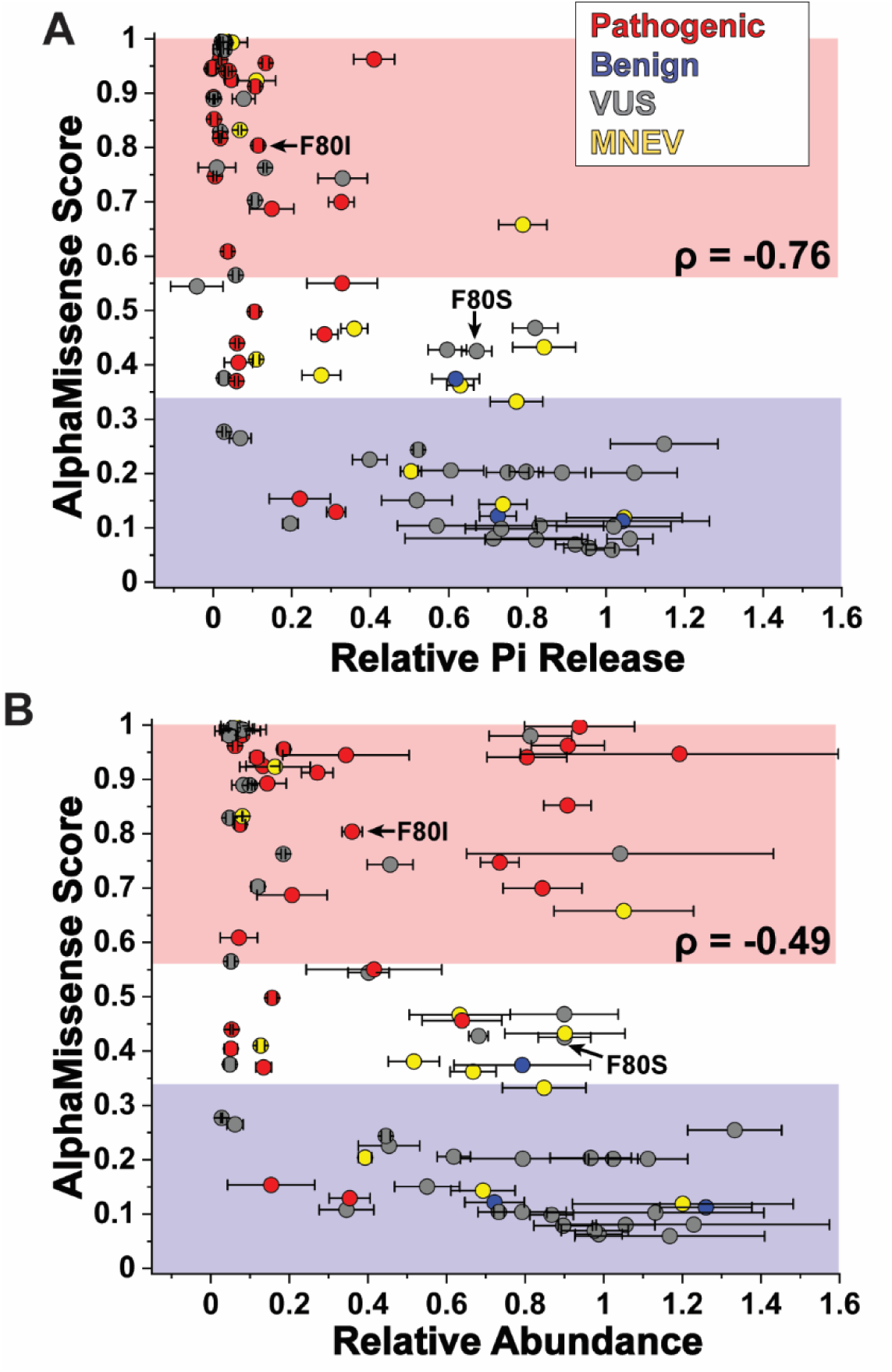
Correlation of AlphaMissense pathogenic probability with experimental metrics. AlphaMissense scores show a stronger correlation with G6PC1 activity (**A**), measured as the Pi released from G6P hydrolysis relative to the WT, than with folded protein abundance (**B**) determined from fluorescence detection size exclusion chromatography of EGFP fusion constructs. As highlighted for two variants with a shared position, F80I/S, the *in vitro* assays discriminate between G6PC1 properties. The standard deviation from the mean derived from n≥3 biological replicates is shown.

Importantly, the analysis could distinguish sidechain-dependent changes in enzyme function for variant positions screened against multiple substitutions, including clinical and non-clinical (MNEV) variants. In most cases, such variants reported attenuated, but not a complete loss of, activity that was in line with shifts in AlphaMissense score. For example, F80I is a GSD1a-linked variant that was scored by AlphaMissense with high pathogenic probability that reported only ∼10% of WT activity (Fig 2A), consistent with the expected disease phenotype. F80S is labeled as “likely pathogenic” in ClinVar, yet much lower in AlphaMissense, EVE and CPT-1 pathogenic probability (Table S1). Inspection of the cited literature only provides genetic evidence for the pathogenicity of F80I (62), and there are no other publications available for F80S in the NCBI databases. We suggest that F80S was erroneously assumed to be disease-causing based on the F80I phenotype. In support of this conclusion, F80S displayed substantially greater activity (6-fold) than F80I, similar to the benign-labeled variant V131A, consonant with BS3 evidence according to recent American College of Medical Genetics standards (9). Informed by our *in vitro* studies here and lack of clear *in silico* evidence and clinical association, we have corrected the F80S classification to “uncertain significance” in Table S2 and all relevant figures.

Interestingly, a weaker correlation was observed in the relationship between AlphaMissense score and variant abundance (Fig 2B). This was caused by variants that displayed disproportionate abundance relative to activity, most of which were found in the “likely pathogenic” regime of AlphaMissense scores. The altered variant profile suggested that changes in G6PC1 abundance are mechanistically linked to disease pathogenesis. Moreover, this result reinforced the conclusion that AlphaMissense predictive accuracy is contingent on both the robustness of the biological metric used to distinguish variant properties (34,38,63) and the underlying mechanism of disruption.

*Mechanistic segregation of variant effects identifies a putative allosteric network*. To further explore the apparent mechanistic divergence more explicitly, we compared directly Pi release of each variant to its abundance relative to the WT (Fig 3A), which uncovered two underlying consequences of the SNVs. First, the activity of most variants corresponded directly to protein abundance, populating an approximate linear relationship (Pearson’s r = 0.78) that sampled the full metric range. We note that only F322Y reported a significant yet proportional increase in both variant abundance and hydrolysis (one-way ANOVA, p<0.0001). We saw no evidence for enhanced hydrolytic capacity (i.e. gain-of-function) where function appeared to exceed variant abundance. In contrast, 20 of 28 GSD1a-linked variants demonstrated ≤ 50% of WT protein levels, indicating that reduced G6PC1 abundance is a major driver of disease pathogenesis.

**Figure 3.**
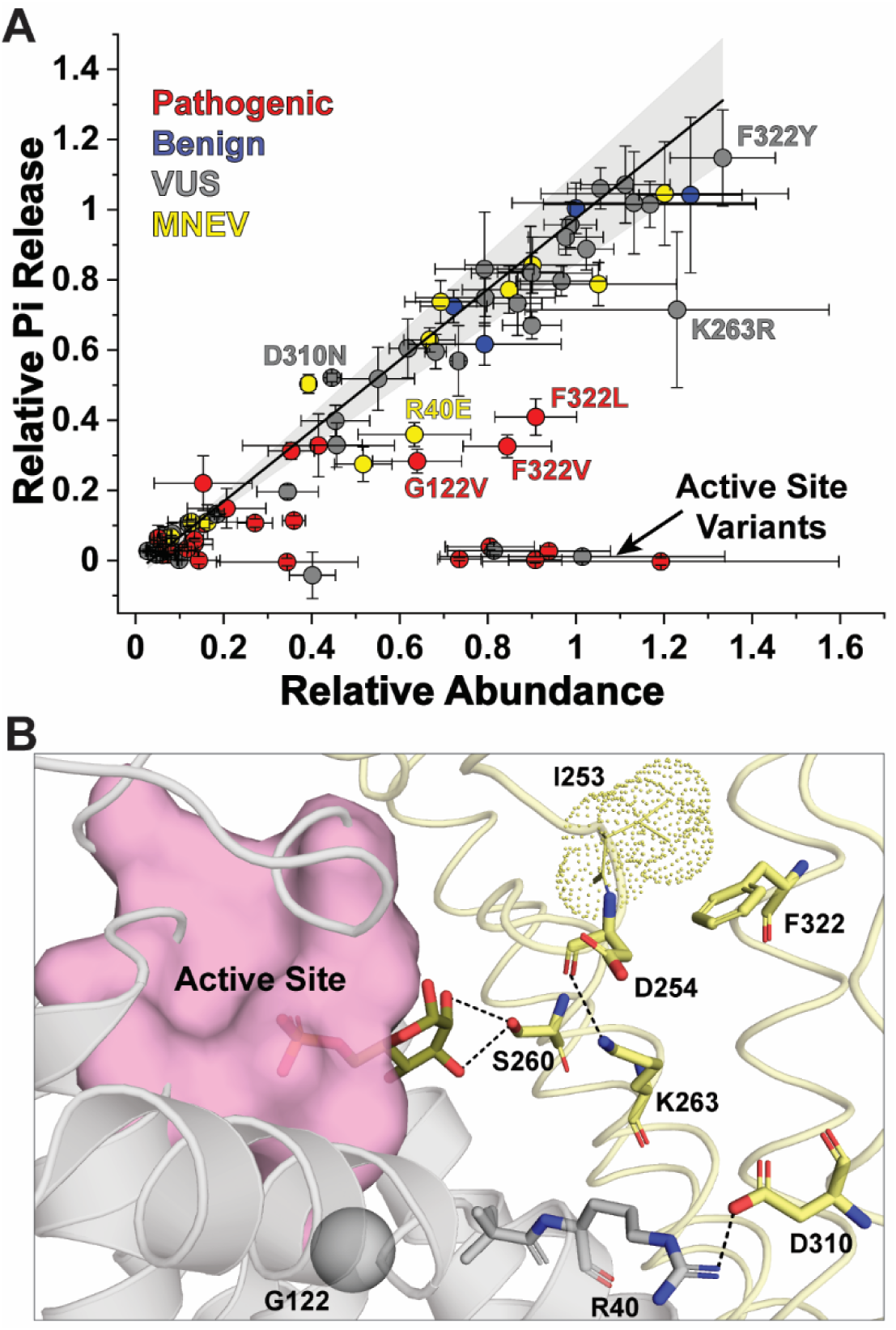
Relationship between folded protein abundance and activity for each variant. **A**. The expression-activity correlation follows a linear regression model (Pearson’s r=0.78, p<0.0001). The solid fit line (slope=1.01) is bounded by a 95% confidence interval (grey). Deviations from linearity are linked to the relative location of the variant within the G6PC1 structure (PDB ID 9JTM), either found in the active site or an adjacent site comprised of largely conserved or homologous sidechains among other G6PC isoforms or vertebrate species (**B**). The PAP2 region is colored grey and the ancillary helices yellow. Hydrogen bonds are indicated by dash lines. G6P is shown in the active site formed by the conserved tripartite sequence motif. The stacking interactions between I253 and F322 is emphasized by the cloud representation of I253.

Secondly, outliers from the linear activity-abundance relationship reported attenuation or the complete absence of activity that did not correspond to folded enzyme abundance. From a structural perspective, these variants were found either exclusively within the active site or as part of a highly conserved and interconnected network of amino acids adjacent to the active site (Fig 3B). We have suggested previously that impaired catalysis in active site variants may be driven by structural distortions and/or disrupted electrostatic interactions (48). Within the PAP2 fold near the active site (Fig 3B), variants of G122 decreased both abundance and catalysis, consistent with pathogenic association (Fig 3A). Despite their location within the ancillary helices (Fig 3B), the sidechains of S260 and K263 project toward the active site in the G6P-bound model and form H-bond interactions with either the G6P hexose ring or with the backbone carbonyl of a highly conserved Asp (D254). Two variants of uncertain significance, S260R and K263E, displayed similar biochemical properties as the pathogenic active site variants (Fig. S2). The SNV K263R largely preserved activity, consistent with a critical role for a positively charged sidechain at this position (Fig 3A).

The broader implications of the conserved residue network were highlighted by the variant profiles of R40, D254, D310 and F322. While perturbed electrostatic interactions (R40/D310) or altered sidechain chemistries (D254) disrupted variant abundance and/or activities disproportionately (Fig 3, Fig. S2), variants at F322 displayed the most divergent biochemical pattern. In stark contrast to the variant of unknown significance F322Y, two other variants, F322L and F322V, demonstrated strongly attenuated function that was consistent with the GSD1a association. As outlined below, the origin of such disparity was likely linked to sidechain repacking and backbone displacement that perturbed functional dynamics. Collectively, the altered activity profiles of variants within this conserved residue network suggest the potential for allosteric modulation of active site hydrolysis. The following sections describe experiments that explored the fundamental bases for the patterns observed in Fig 3A.

*G6PC1 variant abundance correlates with thermostability*. We investigated variant-dependent changes in G6PC1 stability based on the hypothesis that protein stability is deterministic for abundance (64,65). In the absence of explicit measurements of the Gibbs energy of unfolding (ΔG_unf_), the application of protein stability predictors derived from computational resources such as Rosetta, FoldX, or ThermoMPNN, have attempted to estimate changes in ΔG_unf_ (ΔΔG) induced by mutation (66,67). These tools harness either a combination of physical and statistical energy terms (Rosetta (27) and FoldX (28)) or a deep learning approach (ThermoMPNN (68)) as the basis of predictions. Moderate Spearman rank correlations (0.4-0.55 on average) between cellular abundance and predicted ΔΔG have been observed for water soluble proteins (64). However, evaluation of these tools on biochemical metrics or cross correlation with AlphaMissense scores did not yield predictable consequences for G6PC1 variants (Fig. S4). Moreover, the scale of ΔΔG perturbations varied widely between the predictors. The heterogeneous performance of FoldX and Rosetta has been previously noted (23,69–71) as well as sources of bias (72). Accurately predicting ΔΔG for membrane proteins is inherently difficult (73,74) and limited by numerous factors such as imperfectly calibrated score functions, implicit membrane modeling, and dataset composition for training of machine learning models (75,76). For instance, ThermoMPNN did not include membrane proteins in the training dataset, precluding proper inferencing.

Acquisition of thermal denaturation curves has been shown to be a surrogate reporter of ΔG_unf_ with exquisite sensitivity to numerous variables including residue substitution (77,78,60,48). Previously, we demonstrated that thermal tolerance of purified enzyme can be quantified by temperature-induced loss of sample measured by Trp fluorescence detection-size exclusion chromatography to extrapolate a melting temperature, *T*_M_ (60). We adapted the methodology to screen G6PC1 variants within the context of the intact C-terminal EGFP fusion to exploit EGFP fluorescence, which precluded the requirement for purified enzyme. We chose 17 variants that sampled a broad range of protein abundance and activity according to Figure 3. This panel of variants also represented distinct packing environments and sidechain interaction patterns that are electrostatic or hydrophobic in nature. Comparison of the G6PC1-EGFP fusion thermostability curves with corresponding purified samples (devoid of EGFP) indicated that the *T*_M_ differed ∼1 °C on average (Fig. S5), suggesting minimal contribution of EGFP to enzyme stability. While a selection of the results is shown in Figure 4 to emphasize salient trends, the complete dataset can be found in Fig. S5.

**Figure 4.**
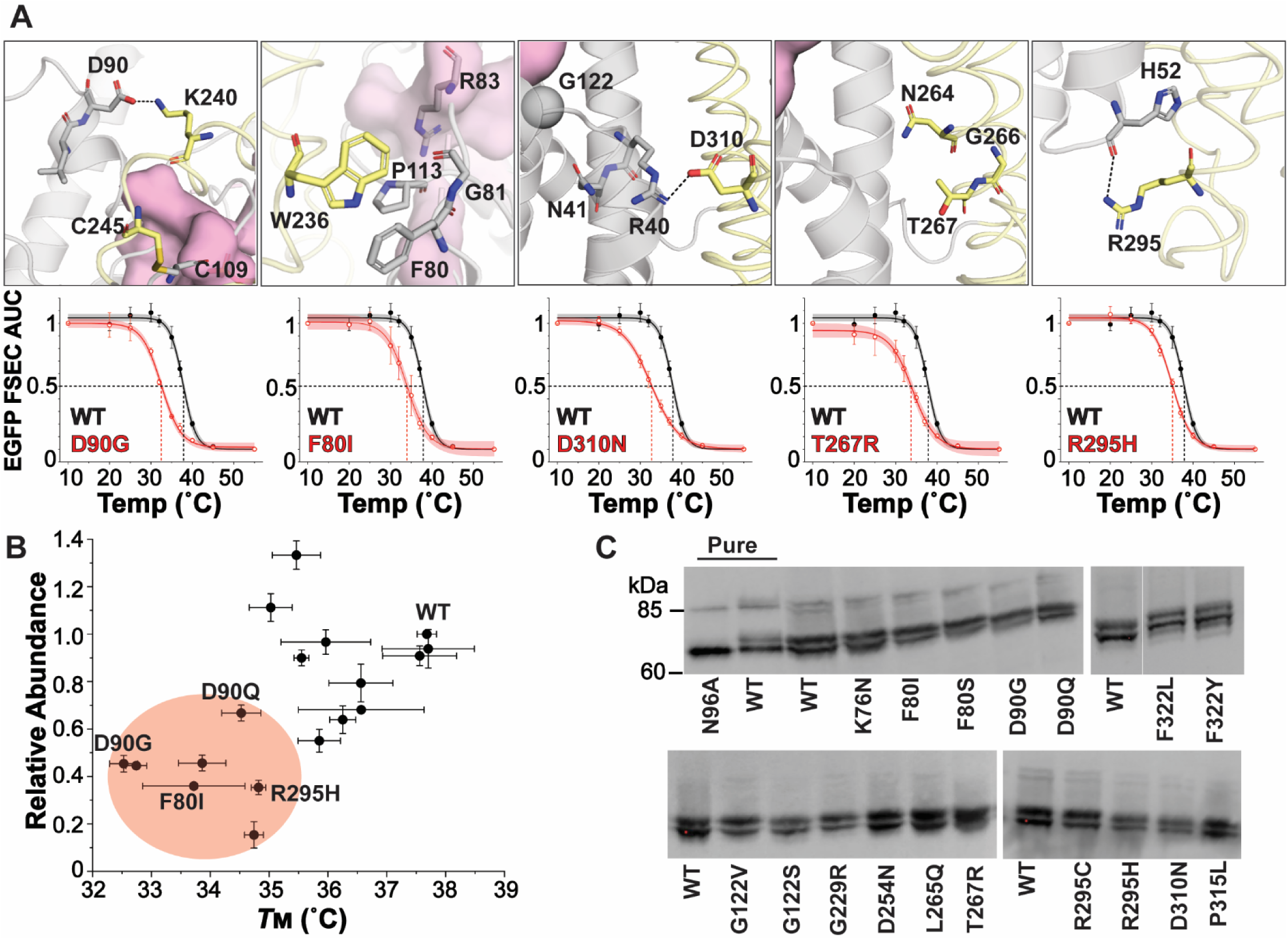
Thermostability profiling of select variants sampling unique structural motifs between the PAP2 fold and ancillary helices. **A.** Location of perturbed sidechain interactions and the corresponding melt curves to define the melting temperature (*T*_M_) from non-linear least squares fitting assuming a dose response model. The standard deviation of each data point was derived from three independent experiments. The fit line is bounded by a 95% confidence interval. **B.** The correlation between *T*_M_ and abundance identifies variants that are susceptible to instability. Those variants with significantly different *T*_M_ relative to WT (one-way ANOVA, p<0.0001) are highlighted by the color-filled region. The data are expressed as mean ± SEM. **C.** Glycosylation of N96 is retained for all variants.

In general, we observed a pattern in which variant abundance not only tracked with its *T*_M_, but also with the chemistry of the packing environment. Figure 4A illustrates the localized structure of select variants and the corresponding thermostability melt curves relative to the WT. These variants demonstrated a statistically significant departure from the WT, reducing the *T*_M_ 3-5 °C, and yielded ≤ 50% of WT protein abundance (Fig 4B). Interestingly, this analysis emphasized the role of packing interactions at the interface of the PAP2 fold and ancillary helices. The epitome of such stabilizing interactions was the disulfide (C109/C245) that covalently links large loops between TM2/3 and TM6/7 and contributes to active site architecture. Substitution of C109 with numerous sidechains, including two clinical variants with H-bonding potential, abolished folded protein expression (Fig. S2). A similar effect of disulfide mutagenesis was reported previously for the pancreatic islet-specific G6PC isoform, G6PC2 (79).

Besides this covalent requirement for folding, disruption of salt bridges that either link the large loops (D90/K240) or anchor the C-terminal region of TM8 to the PAP2 fold (R40/D310) with clinical variants D90G and D310N, respectively, reduced enzyme thermostability by 5°C (Fig 4A). The measured disruption of D90/K240 could be partly mitigated by introducing a polar sidechain (D90Q) that increased abundance by nearly 50% and *T*_M_ by 2 °C relative to D90G (Fig 4B). Accordingly, a reciprocal substitution K240E also reduced abundance and abolished activity (Fig. S2). Though not as severe as D310N, the R295C/H variants perturbed stability of the PAP2/ancillary interface at the opposite end of TM8 likely through loss of Arg sidechain H-bonding with the backbone carbonyl of H52. As expected, the introduction of large positively charged sidechains (T267R) within the transmembrane helix core or the removal of hydrophobic pi stacking interactions (F80I) promoted destabilization of G6PC1.

Despite the limited stability analysis, we found that enzyme abundance for these variants was predictive for the abundance of other variants in the immediate vicinity. Variants at positions highlighted in each panel of Fig 4A displayed <50% of WT abundance (Fig. S2), suggesting that a preponderance of clinical variant expression profiles could reflect regionally destabilized “hot spots” by proxy. To support our interpretation that such substitutions alter intrinsic enzyme stability, we explored glycosylation patterns that should reflect proper enzyme maturation. In a previous study, we discovered that N-linked glycosylation at Asn96 promoted G6PC1 stability without significantly altering hydrolysis kinetics(60). All variants that were screened for thermostability demonstrated a western blot banding pattern consistent with the presence of glycosylated and non-glycosylated enzyme (Fig 4C). Although WT G6PC1 displayed relatively equal populations of both forms, densitometry indicated that several variants were reduced in the non-glycosylated population (Fig. S5), suggesting an increase in either maturation efficiency or, more likely, degradation of non-glycosylated enzyme. Regardless, these results imply that destabilizing variants do not impair G6PC1 N-glycosylation. As a corollary to the *in vitro* thermostability measurements, we investigated whether destabilizing missense variants activated the cellular response to misfolded proteins.

*Destabilized variants activate the unfolded protein response.* The decrease in folded protein abundance because of reduced stability raises the possibility that misfolded G6PC1 stimulates ER stress and activation of the unfolded protein response (UPR). To test this, we generated a plasmid containing multimerized activating transcription factor 4 (ATF4) and activating transcription factor 6 (ATF6) binding sites ligated to a minimal *Xenopus* albumin promoter and the luciferase reporter gene. Both ATF4 and ATF6 are induced by the UPR but combining the binding sites increased sensitivity (Fig. S6). The sequence of the ATF4 binding site (80) and two ATF6 binding sites (81) used in this reporter plasmid had been determined by ChIP-Seq. Figure 5 shows that overexpression of select G6PC1 variants by transient transfection in the islet-derived 832/13 cell line stimulated luciferase expression to different levels, indicating that the strength of UPR activation was variant dependent. Strikingly, the degree of UPR activation was inversely correlated with variant abundance in HEK293SG cells.

**Figure 5.**
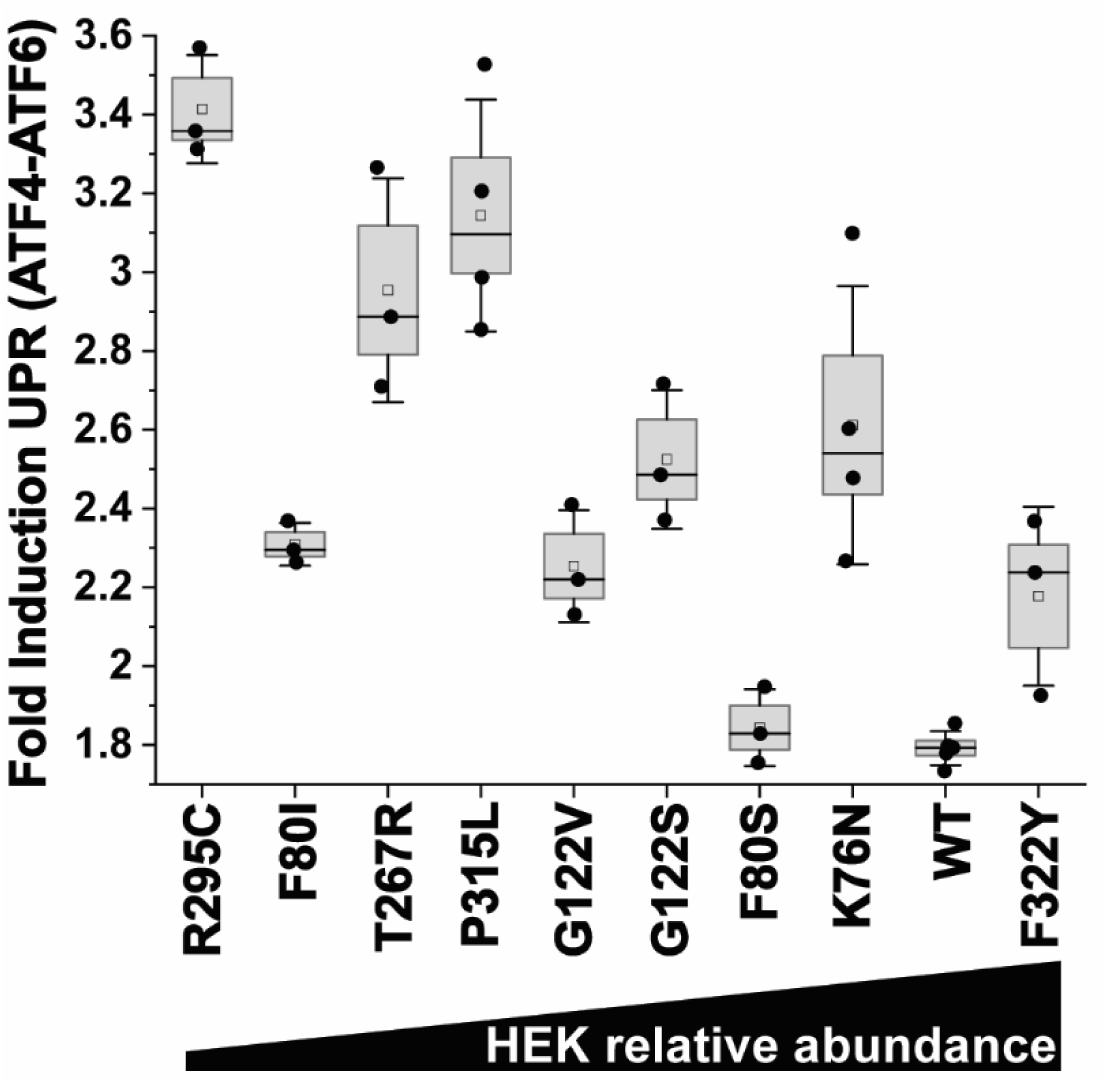
Activation of the unfolded protein response by destabilized missense variants. Induction of luciferase expression by activation of a multimerized ATF4-ATF6 reporter plasmid showed missense variant dependence in the 832/13 islet cell line. Induction strength was inversely correlated with folded enzyme abundance in HEK293SG cells. Measurements were collected from n ≥ 3 experiments, which are shown as individual data points for each variant. The box size reports the standard error and the whisker the standard deviation. The mean is shown as an empty square and the median as a line.

*Site-specific modulation of catalytic properties within the conserved allosteric network*. Despite the unique pattern of biochemical metrics reported by variants in the proposed allosteric residue network (Fig 3), the binary activity profiles cannot distinguish between changes in *K*_M_ or *V*_max_ that alter catalytic efficiency. Steady state kinetic analysis of four purified clinical variants uncovered distinctive properties that reflected the mechanistic modes of allosteric modulation (Fig 6). While Michaelis-Menten fitting parameters of D254N were similar to WT, variants at D310 and F322 demonstrated significant shifts in either *K*_M_ or kcat (Fig 6A,C). D310N demonstrated the most pronounced reduction (35%) in G6P turnover, which could not be attributed to instability at the assay temperature (Fig. S7). However, catalytic efficiency (kcat/*K*_M_) remained at WT levels due to compensatory changes in *K*_M_ (Fig 6C).

**Figure 6.**
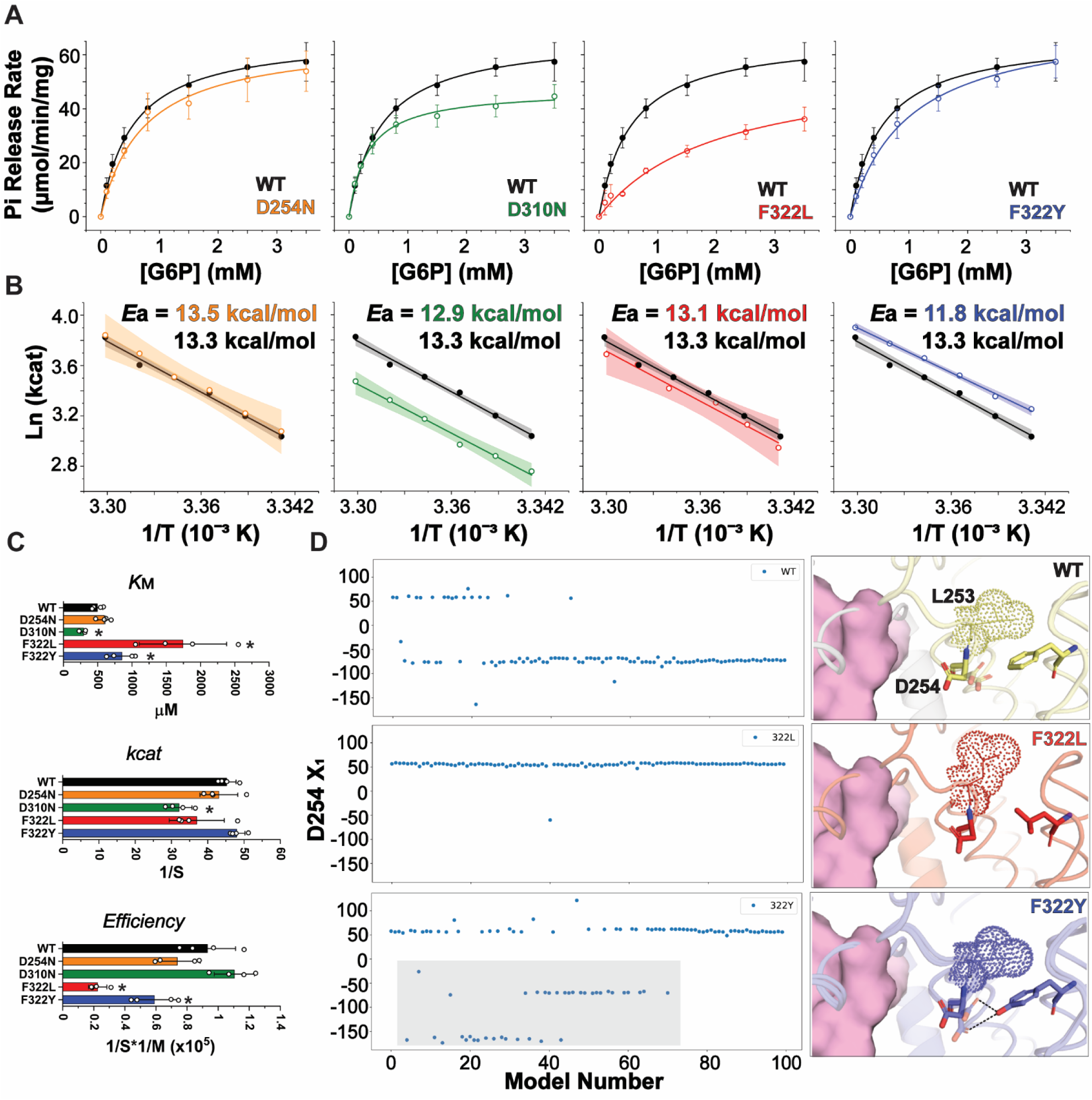
Kinetic, thermodynamic, and structural modeling of variants in the allosteric network. **A**. Steady-state kinetics of G6P hydrolysis mediated by purified protein at 30 °C illustrates distinct activity profiles. **B.** The temperature dependence of hydrolysis indicated that the F322Y variant lowers transition state energy by ∼1.5 kcal/mol, consistent with the strength of a hydrogen bond. **C.** Altered kinetic properties highlight changes in the Michaelis constant and catalytic efficiency. Asterisks (*) denote statistical significance relative to the WT in a one sample t test. For *K*_M_ of D310N and F322 variants, p = 0.003-0.04. For kcat of D310N, p = 0.003. For catalytic efficiency of F322 variants, p = 0.00016-0.02. **D.** AlphaFold modeling of WT, F322L and F322Y suggests changes to hydrogen bonding potential based on an altered rotameric distribution of D254, rationalizing the changes in activation energy. The grey box identifies D254 rotamers capable of hydrogen bonds with Y322.

The stark differences in variant profiles at residue 322 captured by the binary assay (Fig 3) were linked to perturbations in both kinetic and thermodynamic parameters. In both F322L and F322Y, *K*_M_ was shifted significantly, yet unequally, relative to WT leading to a substantial decrease in catalytic efficiency. Interestingly, the activation energy (*E*_a_) of F322Y, which is defined by the slope of the linearly transformed relationship between kcat and temperature, was clearly divergent from the WT (Fig 6B). The difference in *E*_a_ (1.5 kcal/mol on average) was approximately the strength of a weak H-bond. In the cryo-EM structure of the G6P-bound state (82,83), F322 was found to make hydrophobic contacts with I253 (L253 in the mouse ortholog), which likely supports active site closure mediated by the large loop between TM 6 and 7 (Fig. S8). This interaction is absent without substrate occupation of the active site due to a distinct conformation of the TM6/7 loop (Fig. S8). Since substitution of F322 with smaller branched amino acids could compromise packing with residue 253, we suspect that the hydroxyl moiety of Y322 could form H-bond interactions with the D254 sidechain to stabilize a “closed loop” conformation of G6PC1 that mimics the G6P-bound state. This hypothesis would rationalize the differences in catalytic efficiency between F322Y and F322L assuming that loop closure correlates with G6P hydrolysis.

Since AlphaFold models have been shown to be sensitive to local structural deformations caused by amino acid substitutions (84,85), we investigated if AlphaFold could predict nuances of sidechain and backbone packing for F322 variants (see Methods for details). Interestingly, AlphaFold modeling of L322 and Y322 provided support for our hypotheses, illustrating potential displacements and alternative rotamers of D254 within an ensemble of 100 predicted structures (Fig 6D, Fig. S8). In particular, Y322 increased the predicted population of a D254 rotamer that was observed in only one WT model (χ≍-150°) with potential for H-bonding to the hydroxyl moiety of Y322. As a first-order approximation, these results implicate variant-dependent shifts in sidechain and/or backbone conformation that alter catalytic properties.

## Discussion

The global analysis of G6PC1 variants described here emphasized two fundamental pathways perturbed by missense variants, namely protein stability and allostery. The activity of most variants followed a simple relationship with abundance, i.e. total Pi release from G6P hydrolysis corresponded to the abundance of folded G6PC1. While *in vitro* thermal tolerance assays linked abundance to intrinsic G6PC1 stability (*T*_M_), the *in situ* consequence of reduced stability was activation of the UPR. Given that ER stress and UPR activation are observed in a number of liver pathologies (86), we speculate that ER stress stemming from destabilized G6PC1 variants contributes to the GSD1a phenotype. Moreover, our results resonate with a recent large-scale variant analysis of many proteins projecting that ∼60% of pathogenic mutations are associated with altered stability (19). We observed that 20/28 ClinVar “pathogenic” variants displayed < 50% of WT abundance, and we predict these variants are likely destabilizing. More broadly, 43 variants at 32 residue positions showed strongly attenuated enzyme abundance. In line with prior studies from the Chou laboratory (47), the vast majority of these are found in the transmembrane helices. In contrast with the Chou study (47), we also found that variants within structured or unstructured loops near the active site compromise protein abundance. The origin of this discrepancy could be differences in the enzyme detection scheme. Whereas the Chou laboratory employed western blotting that can be prone to non-linearities, our chromatographic approach differentiates folded from misfolded and/or aggregated protein. These results imply that therapeutics designed to promote stabilization of the folded state is likely to be beneficial, similar to the strategies for certain destabilizing CFTR missense variants (87). However, we note that the present analysis did not include other factors that can underly enzyme abundance, such as mRNA stability (88–90), translation efficiency (91,92), or degron changes (93). These unexplored factors may have contributed to the moderate correlation between abundance and *T*_M_ (Spearman’s RCC=0.62, p=0.00582).

Variants that retained >50% WT-like abundance yet with consequences to G6PC1 hydrolysis kinetics were localized to a relatively conserved network of sidechains adjacent to the active site and mostly found in the ancillary helices (Fig 3B). There are notable differences in sidechain identity among the three G6PC isoforms, namely at residues 40, 260, 263 and 310 (G6PC1 numbering). In contrast, D254 and F322 are strongly conserved (Table S1). Our results suggest that this residue network may contribute to distinct kinetic properties between isoforms. Supporting this conclusion, G6PC1 activity is ∼80x higher than G6PC2 under similar experimental conditions (79). Within the context of the cryo-EM structures (82,83), this region is dynamic and is expected to regulate substrate access to the active site (Fig. S8). Interestingly, non-protein electron density attributed as a lipid and presumed to be POPS has been modeled in this region with H-bond and electrostatic interactions between D38, R40 and D254 and the lipid headgroup (83). Though a mechanistic role for this lipid remains to be determined, Chen et al proposed that POPS binding could stabilize a catalytically competent conformation. Collectively, we propose that G6P hydrolysis can be allosterically modulated by an altered pattern of sidechain and/or lipid interactions that shift protein conformation. Though high affinity competitive inhibitors of G6PC1 are prohibitive physiologically (94), this model implies that rationally designed small molecules targeting this allosteric residue network may be a viable therapeutic avenue for treatment of GSD1a.

Including active site variants studied previously, patterns arising from the collective dataset of 85 variants further stimulate hypotheses with clinical implications. Remarkably, the divergent distribution of relative activities for known pathogenic and benign variants (Fig 2A and Fig 3A) potentially suggests a threshold for the GSD1a phenotype, facilitating molecular classification based on intrinsic activity. Demarcated by F322L (ClinVar “likely pathogenic”, relative activity of 0.41±0.05) and V131A (ClinVar “likely benign”, relative activity 0.62±0.06), we conservatively assign the variants tested here as likely pathogenic if their relative *in vitro* activity is < 50% of WT. Supporting this conclusion, individuals heterozygous for debilitating G6PC1 variants escape the GSD1a phenotype presumably through compensation by the functional allele (95). The Sankey diagram in Figure 7A captures the classification trail based on ClinVar, AlphaMissense and our experiments. From this activity threshold, we find that AlphaMissense accurately classified 90% of predicted benign and pathogenic variants (AUROC = 0.93, p<0.0001).

**Figure 7.**
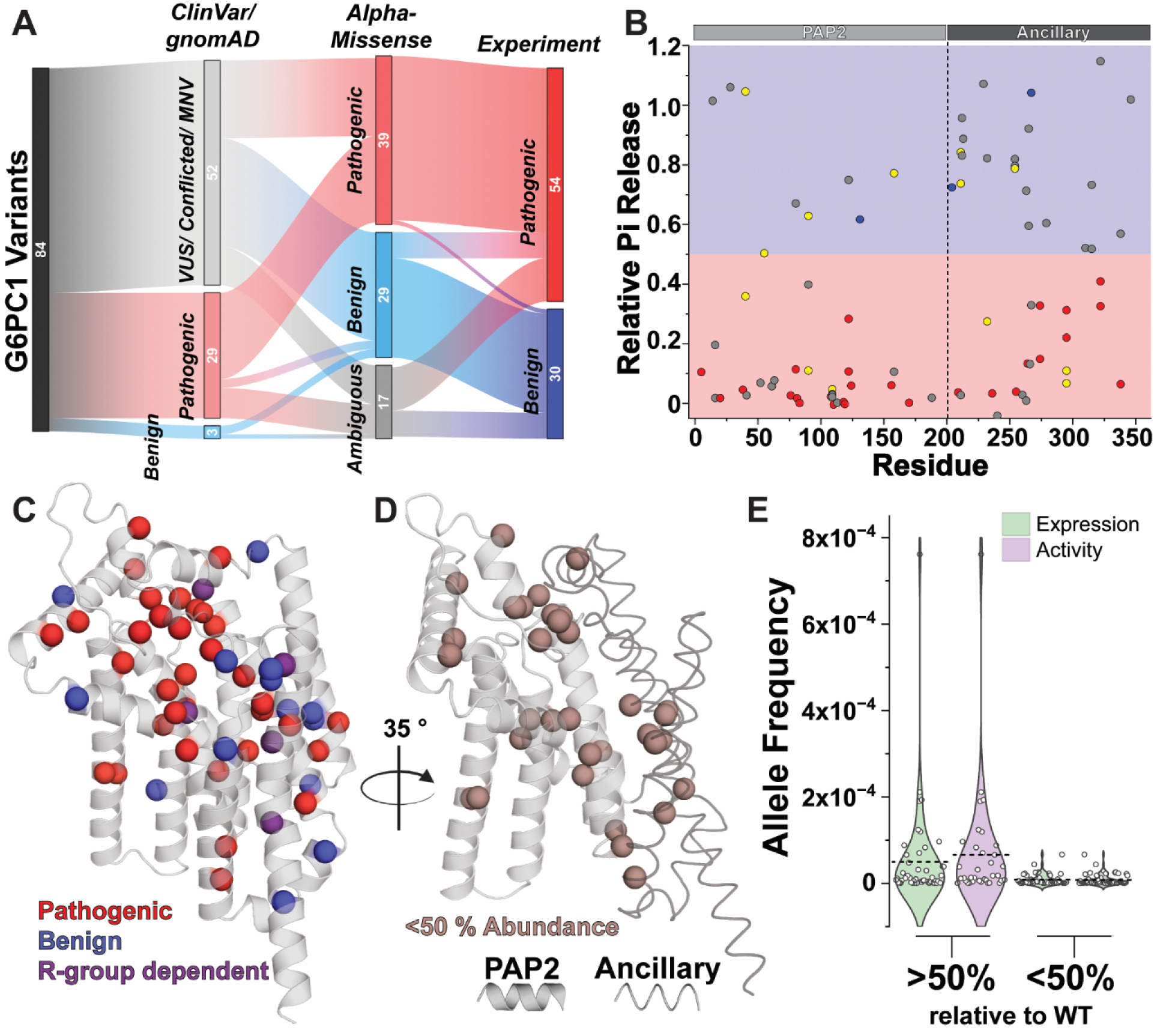
Classification of molecular phenotypes and the principles of G6PC1 variant interpretation. **A.** Flow chart of clinical classifications based on distinct levels of variant review. **B.** The number of variants characterized arising from the PAP2 fold and ancillary helices were approximately equal (43 at 29 residue positions and 42 at 25 residue positions, respectively). The relative density of pathogenic variants is greater in the PAP2 fold, which is a highly conserved feature of the PAP2 superfamily. **C.** The tertiary distribution of experimentally classified variants emphasizes the difficulty of pathogenicity prediction based on structure alone. **D.** Variants that perturb the interface between the PAP2 fold and ancillary helices induce instability. **E.** Violin plots display the distribution of alternate allele frequencies based on an empirical threshold for variant abundance and activity. Alternate allele frequency cannot discriminate molecular phenotypes interpreted from experimental metrics. The dashed line represents the mean allele frequency.

Although our analysis is limited to ∼10% of all observed SNVs, principles for variant interpretation can be derived from trends observed in the broad variant panel. As the defining feature of the PAP2 superfamily, the PAP2 fold appears more susceptible to disruptive amino acid variation than those in the ancillary helices (Fig 7B). Except for the pathogenic “hot spot” within the active site, a structure-only prediction of disease probability is not likely feasible based on the wide distribution of benign and pathogenic sites (Fig 7C). Nevertheless, assuming abundance as a proxy for stability, the pattern of variants with strongly attenuated expression highlights the importance of stabilizing packing interactions at the PAP2/ancillary interface (Fig 7D). As discussed elsewhere (52,96–98), we emphasize that simply knowing the alternate allele frequencies for these very rare variants is not sufficient for pathogenicity prediction. We parsed allele frequencies derived from the gnomAD_v4.1 and RGC databases based on our protein abundance or activity thresholds (Fig 7E). Even though variants with benign molecular phenotypes tend to be observed more often on average, pathogenic variants were found to have similar frequencies albeit with a tighter distribution.

To our knowledge, this collective body of work represents the most comprehensive biochemical analysis of missense and functional variants in G6PC1 to date and is consistent with results from other investigations (51,82,83). Although this study has uncovered mechanistic determinants of catalysis and dysfunction in a recessive Mendelian disorder, the significance of these results could be markedly bolstered by leveraging genotype-phenotype association studies from biobank datasets. In particular, the hypothesis of a pathogenic threshold necessitates an extensive analysis of variant gene expression coupled with assays of protein abundance and catalytic efficiency to define the “benign” phenotype and establish the activity gradient that precludes disease. Such studies would provide a benchmark for therapeutics designed to recover adequate G6PC1 function in patients with GSD1a.

## Conclusions

The premise of our investigation asserts that patterns arising from population genetics and sequence evolution augmented by state-of-the-art, structure-aware pathogenicity prediction can guide targeted mechanistic inquiry of missense variants through discovery of molecular “fingerprints” that rationalize the elementary disease phenotype. We exploited AlphaMissense pathogenicity predictions to support selection of a relatively sparse panel of G6PC1 variants with unique clinical classifications that broadly sample sequence and structure space. While our analysis identified erroneous predictions, the strong correlation between AlphaMissense score and G6PC1 variant activity suggests that the tool can stimulate hypothesis-driven studies that bridge the mechanistic divide between genotypes and phenotypes provided that experimental metrics are directly linked to target protein properties. Moreover, our results imply that even minimally stratified datasets (multiple substitutions per position) can deliver rich information content when coupled with sensitive assays that report unambiguous molecular phenotypes. If visualized as a strategic continuum bounded by computational prediction and saturation mutagenesis extremes, this approach represents an intermediary that borrows the evolutionary information encoded within a protein language model and the gene-specific experimental properties to capture salient mechanistic features linked to missense variation. We suggest that this approach will be advantageous for characterizing targets which lack broad biochemical foundations to interpret missense variant phenotypes without over-reliance on strict *in silico* predictions or expensive deep mutational scanning assays.

## Methods

### Bioinformatic analysis and AlphaMissense predictions

Table S1 summarizes the attributes of 549 non-synonymous SNVs manually curated from four databases, namely ClinVar, the Human Gene Mutation Database (HGMD), gnomAD_v4.1, and the Regeneron Genetics Center Million Exome Variant Browser (RGC). Allele frequencies for 413 and 347 variants were pulled from gnomAD_v4.1 and the RGC, respectively. The RGC database included allele frequencies for 80 unique variants relative to gnomAD_v4.1, yielding a total of 493 variants with an allele frequency label. As a whole, the allele frequencies of the 267 variants common to both databases were not statistically different (unpaired t test, p=0.5281) and 245 of these variants (92%) differed by less than a factor of 2. To represent the combined allele frequency from both databases, we created a simple “substitution index” by aggregating the allele frequencies for each variant per residue from gnomAD_v4.1 and merging the data of unique variants from the RGC group. For example, SNVs of Phe80 generate Ile (6.21E-7), Tyr (1.86E-6), Ser (4.35E-6), and Val (6.08E-7) substitutions with a substitution index (sum) of 7.44E-6. The substitution index thus represents the observed frequency of variation per residue, which was visualized using ChimeraX (99) molecular viewer in Fig 1a.

The evolutionary conservation of the G6PC1 primary sequence was estimated by ConSurf (49,50) initiated with a multiple sequence alignment composed of 1000 sequences derived from an MMseq2 search of UniRef100. The phylogenetic tree was reconstructed using a neighbor-joining algorithm with an interpretation assuming the JTT evolutionary model and the site-specific evolutionary rate (conservation scores) determined from an empirical Bayesian paradigm. Table S1 reports the most common amino acid, its relative population percentage, and the conservation score for each residue observed to undergo single nucleotide variation. The conservation score informs the color scale of Fig 1d.

AlphaMissense scores can be obtained from the Ensembl (100) genome browser, as a large file on Zenodo (101), or from a searchable web resource (102). Table S1 and S2 report the AlphaMissense (AM) scores and class labels for the 549 observed SNVs and our tested targets, respectively.

### G6PC1 construct design and mutagenesis

Mouse G6PC1 (mG6PC1, UniProt P35576) was used as a model WT template for introduction of amino acid substitutions into the primary sequence. Relative to human G6PC1 (hG6PC1, UniProt P35575), mG6PC1 retains 89% sequence identity as well as a conserved predicted tertiary architecture, and is subject to the same regulation *in vivo* (59). However, we have shown that mG6PC1 has a greater inherent stability *in vitro* than hG6PC1 (60), effectively increasing the dynamic range (sensitivity) of biochemical and biophysical measurements. We refer to mG6PC1 as “G6PC1” for simplicity throughout the manuscript. Similar to our previously published work (48,60), G6PC1 was expressed as a fusion protein with the sequence of enhanced green fluorescent protein (EGFP) followed by a poly-histidine (His_8_) tag on the C-terminus. The fusion construct was cloned into the pJPA5 vector 3’ of the CMV promoter for expression in mammalian cells. Substitutions in G6PC1 were generated using site-directed mutagenesis and amplified through PCR protocols using the Q5 polymerase (New England Biolabs). Bacterial transformation was performed with Dpn1-digested PCR products into E. coli XL1 competent cells (Vanderbilt University Cell Culture Core). The resulting DNA plasmid was purified with the Qiagen Spin Miniprep Kit (Qiagen). Full DNA sequencing of the G6PC1 insert verified desired substitutions and absence of unwanted changes.

### Screening of G6PC1 variant expression and activity

G6PC1 variants were introduced into adherent human embryonic kidney (HEK293SG) cells (N-acetylglucosaminyl-transferase I-negative; ATCC CRL-3022) cultured in DMEM:F12 medium supplemented with 10% FBS with Lipofectamine 2000 reagent (Invitrogen) and OptiMEM I Reduced Serum Medium (ThermoFisher). Cells were transfected using 2 μg of DNA per well (two wells per variant) in 6-well plates for 48 hours at 37°C with 7% CO2. Transfection of WT mG6PC1 was used as a positive control for each experiment and a normalizing factor for abundance and activity. Variant expression was confirmed by cell EGFP epifluorescence on either a Zeiss Axio Zoom.V16 fluorescence stereoscope or a Bio-Rad Fluorescence Cell Imager.

Cells were harvested, washed, and solubilized using a previously described protocol (60). Briefly, whole cells were solubilized in 50mM Tris pH 8, 150mM NaCl, 1mM EDTA buffer supplemented with 5mM PMSF and 5mM lauryl maltose neopentyl glycol (LMNG) detergent for 1hr at 4 °C with gentle nutation. Insoluble material was removed by centrifugation at 105000 rcf for 30 minutes in an Optima MAX XP ultracentrifuge. Supernatant containing solubilized G6PC1 was injected onto a Superose6 Increase 10/300 GL size exclusion chromatography column (Cytiva) equilibrated in 50mM Tris pH 8.0, 150mM NaCl, 1mM EDTA, 0.2mM LMNG buffer to determine sample homogeneity and quantify abundance. The column was attached to a Shimadzu LC40i high-performance liquid chromatography system equipped with a fluorescence detector and temperature-controlled multi-sampler. Variant abundance was quantified from the area under the elution curve monitored by EGFP fluorescence (ex 475nm, em 515nm) and normalized to the WT signal. Variant abundance profiles (Fig. S2) were obtained from the average and standard deviation of at least three biological replicates.

G6PC1 variant activity was characterized by an established end-point colorimetric assay that quantifies the release of Pi from sugar phosphates (60). Briefly, supernatant containing detergent-solubilized G6PC1 was diluted into 50mM Tris/Mes pH 6.5, 50mM NaCl, 0.2mM LMNG buffer in the presence or absence of 1.5mM G6P on ice. Hydrolysis was stimulated by transferring the reaction to a 30°C water bath for 5 minutes followed by quenching with a 2-minute ice incubation and then addition of 12% (w/v) SDS. Color was developed as previously described (60) and the reactions transferred to a clear 96-well microplate for measuring sample absorbance at 850nm on a BioTek Synergy H4 microplate reader. A standard Pi curve was used to determine the amount of Pi released. Each variant was tested in triplicate per biological replicate (n ≥ 3).

### Thermostability profiling of G6PC1 variants

Transfected HEK293SG cells were harvested and solubilized with 1100 μL of 5mM LMNG buffer supplemented with PMSF as described above. Following extraction and ultracentrifugation, 110 μL of supernatant was distributed into PCR tubes for thermocycler incubation at different temperatures: 10 °C, 20 °C, 25 °C, 30 °C, 32 °C, 35 °C, 37 °C, 40 °C, 45 °C, 55 °C. The sample was heated for 10 minutes at its designated temperature. The samples were then subjected to ultracentrifugation at 105000 rcf for 15 minutes to remove precipitated material. The supernatant was injected onto a Superose6 Increase 10/300 GL size exclusion chromatography column (Cytiva) monitoring the EGFP signal (ex 475nm, em 515nm). The area under the elution curve was plotted against the corresponding temperature and fit with a dose response model in Origin (OriginLab) to determine the melting temperature *T*_M_. The mean and standard deviation for each temperature was quantified from three biological replicates. The 95% confidence interval of the fit is shown in Fig 4 and Fig. S5. The *T*_M_ acquired from non-purified samples was compared to the *T*_M_ from purified samples devoid of the EGFP-His_8_ tag (described below) using a similar protocol but monitoring Trp fluorescence (ex 295nm, em 320nm).

### Detection of glycosylation patterns via western blotting

The concentration of G6PC1 variants solubilized from whole HEK293SG cells was estimated from the area under the curve obtained from EGFP fluorescence detection size exclusion chromatography against a purified EGFP standard curve. Approximately 16 ng of each G6PC1 variant was loaded with 0.1M DTT and 1.5x SDS dye into an Invitrogen NuPAGE 10% Bis-Tris gel and electrophoresed at 200V for 1.5 hours on ice using NuPAGE MOPS SDS Running Buffer. The samples were transferred to a Bio-Rad 0.45uM nitrocellulose membrane as described by the Invitrogen Mini Blot Module using NuPAGE Transfer Buffer with 10% (v/v) methanol. The blot was then washed with TBS-T (20mM Tris, 150mM NaCl, 0.1% Tween) and blocked with 5% skim milk in TBS-T by rocking at room temperature for 1 hr. Primary antibody was rabbit anti-green fluorescent protein (Invitrogen #A-11122) at 1:20,000 for overnight probing at 4 °C, and secondary antibody was donkey anti-rabbit IgG (IRDye 680LT LI-COR #92668023) at 1:15,000 for probing at 4 °C for 45 minutes. G6PC1 glycosylated and non-glycosylated bands were profiled by pixels with Fiji (103) and quantified by area under the curve assuming a two-gaussian model in Origin (OriginLab) to determine the relative ratio.

### Unfolded protein response reporter assay

The construction of a rat G6pc1-luciferase fusion gene containing promoter sequence between –7248 and +62 in the pGL3 MOD luciferase vector has been previously described (104). A fusion gene containing multimerized activating transcription factor 4 (ATF4) and activating transcription factor 6 (ATF6) binding sites ligated to a minimal Xenopus albumin promoter and the luciferase reporter gene was generated as follows: The use of a minimal Xenopus albumin promoter to study insulin-regulated gene transcription in the context of the XMB CAT plasmid has been previously described (105). Oligonucleotides representing this promoter were ligated into the HinD III – Xho I digested pGL3 MOD luciferase vector (106) as previously described (107). Multiple copies of the following oligonucleotide containing an ATF4 binding site (80) (underlined) and two distinct ATF6 binding sites (81) (in bold) were then ligated into the HinD III site of the albumin-luciferase fusion gene:

Sense: 5’-AGCT**ACCACGTGG**GATC<u>ATGATGCAAT</u>GATC**AGCCAATCGG**

Antisense: 5’-AGCT**CCGATTGGCT**GATC<u>ATTGCATCAT</u>GATC**CCACGTGGT**

UPR activation was assayed by co-transfecting semi-confluent 832/13 cells in 3.5 cm diameter dishes with 2 μg of the ATF4/ATF6 Xenopus-luciferase plasmid, 0.5 μg of SV40-Renilla luciferase plasmid and 1.0 μg of a mouse G6PC1 expression vector, encoding wild type or mutant protein as indicated, or the empty pJPA5 vector, using the lipofectamine reagent (InVitrogen, Waltham, MA) as previously described (104). Following transfection, cells were incubated for 18-20 hours in serum-free medium supplemented with 30 mM glucose. Cells were then harvested using passive lysis buffer (Promega, Madison, WI). Firefly and Renilla luciferase activity were assayed using the Dual Luciferase Assay kit (Promega, Madison, WI). To correct for variations in transfection efficiency, the results were calculated as a ratio of firefly to Renilla luciferase activity. Results are presented relative to the ratio obtained in the presence of the empty pJPA5 vector. Fusion gene expression was assessed in multiple transfections (n≥3).

### Recombinant expression and purification of G6PC1 variants

G6PC1 was expressed and purified from Sf9 insect cells using established procedures (60). Briefly, the G6PC1-EGFP-His_8_ fusion construct was subcloned into pFastBac1 for baculovirus generation and transduction into Sf9 insect cells. The enzyme was extracted from the membrane fraction by solubilization with 5 mM LMNG in 50mM Tris pH 8, 100mM NaCl, 10% (v/v) glycerol for 1 hr on ice. Insoluble material was removed by ultracentrifugation at 185000 rcf for 1 hr. The supernatant was filtered with a 0.45-μm syringe filter and mixed with Ni^2+^-NTA Superflow resin (Qiagen) in the presence of 25mM imidazole for 4 hr at 4 °C with gentle shaking. The resin was washed with 10 column volumes of buffer containing 100mM imidazole and eluted with 300mM imidazole. The EGFP-His_8_ tag was cleaved by thrombin at 15 °C for 16 hr. The final sample was prepared by gel filtration through a Superose6 Increase 10/300 GL column (Cytiva) equilibrated with 50mM Tris pH 7.5, 150mM NaCl, 0.2mM LMNG. G6PC1 concentration was determined from absorbance reading at 280nm assuming an *A*^0.1%^=2.18 mg/mL. Sample purity was assessed by tryptophan fluorescence detection size exclusion chromatography and SDS-PAGE.

### Steady-state kinetic analysis of purified G6PC1

The rate of Pi release was determined as previously described (60) and shown in Figure 6. G6PC1 variants (0.1 μg) was titrated with increasing concentrations of glucose-6-phosphate in a 30 °C water bath for 1 min and then quenched with 12% (v/v) SDS. The reaction was developed for color and extrapolated to the amount of Pi present using a standard curve. The resulting hydrolysis hyperbola was fit assuming a single site Michaelis-Menten model in Origin (OriginLab) to quantify *K*_M_, *V*_max_, and kcat. The reported kinetic parameters in Fig 6 were derived from four independent curves. The activation energy (*E*_a_) was determined from the slope of the linear fit line describing the relationship between kcat and temperature. For this analysis, kcat was determined from full Michaelis-Menten curves measured by two independent experiments at 20-30 °C in 2 °C intervals. A 95% confidence band was included to show the uncertainty in the regression.

### AlphaFold modeling

An initial multiple sequencing alignment (MSA) for mG6PC1 (UniProt P35576) was generated with MMseq2 utilizing Colabfold. For each variant the corresponding residue changes were made in the equivalent amino acid position across all sequences, following the principles for modeling alternate conformations with AlphaFold2 (85). These modified MSAs were then processed with colabfold using all 5 models with 20 replicates each for 100 models total and ranked using pLDDT.

### Statistics

Unless otherwise stated, all replicated data are expressed as the mean with standard deviation derived from at least three biological iterations. Statistical significance was assessed by either a one sample t test for comparisons between two samples, or a one-way ANOVA followed by a post hoc Tukey for multiple comparisons. Continuous p values are reported where possible; otherwise, the standard convention of p<0.05 was assumed for statistical significance. Statistical analyses were performed in Origin software (OriginLab).

### Data availability

All data is made available in the manuscript and supplementary material.

### Author contributions

R.A.S, E.M.H, R.M.O. and D.P.C. contributed to experimental design, execution and data collection. R.A.S. and D.P.C. identified G6PC1 variants for analysis. R.A.S. and D.P.C. performed variant effect predictions and modeling studies with AlphaFold. E.M.H, C.M.N., M.H.R., and S.A.H. performed molecular biology, G6PC1 abundance and activity assays, and thermostability analysis. D.P.C. G6PC1 purification and thermostability measurements with assistance from T.S. R.M.O. designed the UPR assays and A.M.L. and V.R.D.S performed the UPR assays. R.A.S. and D.P.C. interpreted the data. D.P.C. drafted the manuscript with assistant from R.A.S for figure generation. R.A.S, R.M.O, and D.P.C. edited and revised the manuscript. All authors approved the final version of the manuscript. In the early stages of this work R.A.S and E.M.H contributed in different, but equal aspects of the project. R.A.S was assigned lead author due to a continued role in the project and assistance with manuscript editing and revision.

## Acknowledgements

This work was supported by National Institutes of General Medical Sciences (NIGMS) grant R01GM149686 to D.P.C. We thank Mr. Alec S. Rodriguez for support with molecular biology and assistance with the UPR assays.

## Supplementary Information for

**Figure S1.**
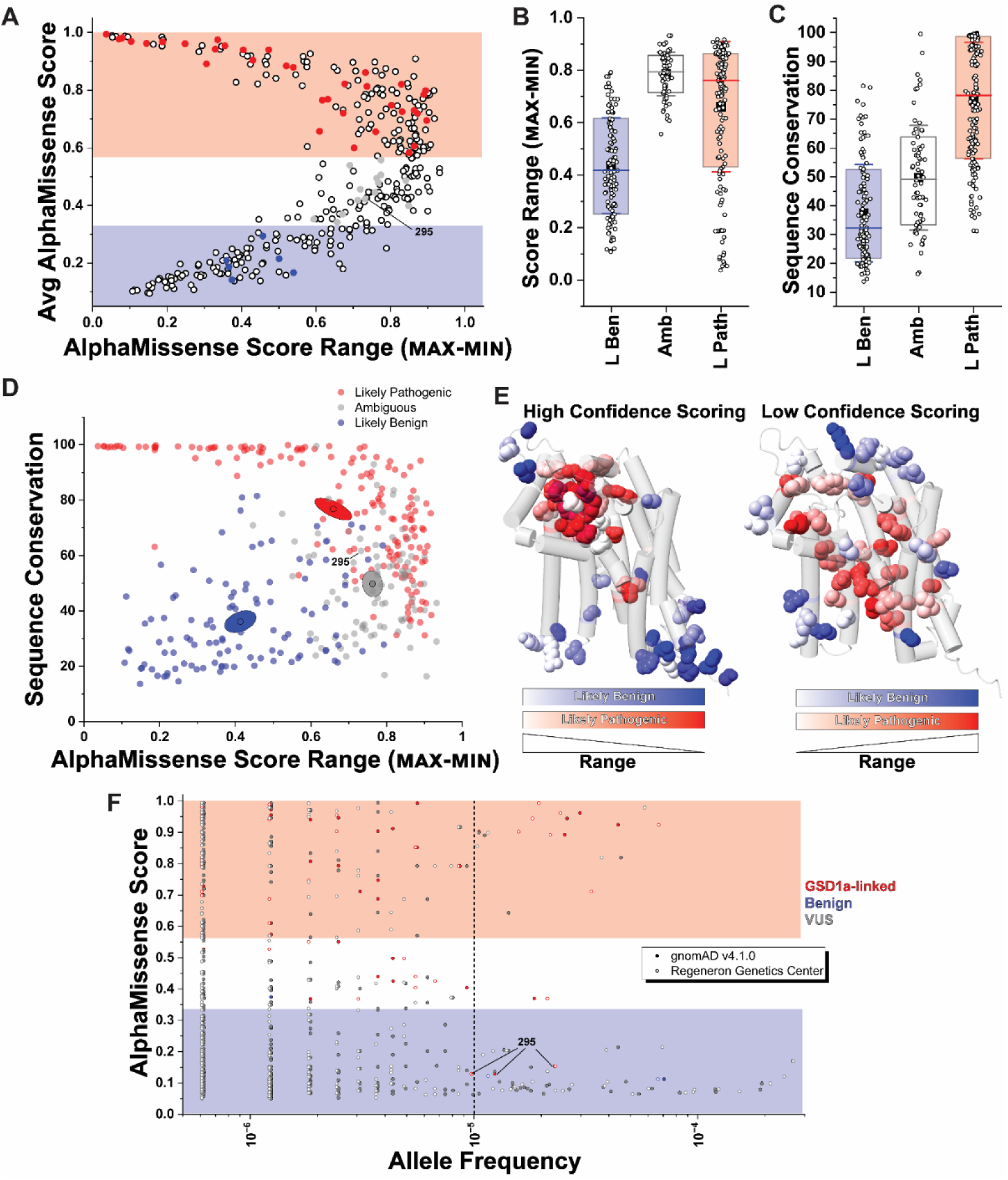
Decomposition of AlphaMissense scoring components. **A.** The average AlphaMissense score for each amino acid in G6PC1 plotted as a function of the AlphaMissense score spread. The solid circles represent the variants examined in this study. **B.** In contrast to the “likely benign” and “likely pathogenic” class, residues in the “ambiguous” class have highly variable AlphaMissense scores across 19 amino acid substitutions and demonstrate intermediate levels of sequence conservation estimated from ConSurf **(C)**. In both **B** and **C**, the boxes encompass 80% of the data, the whisker represents the standard deviation, the mean is represented by a filled black box and the lines identify the median. **D.** The correlation between sequence conservation and AlphaMissense score range highlights overlapping AlphaMissense classifications. Centroids for each class are shown, which captures the average attributes for the given class. **E.** Mapping of the AlphaMissense score range onto a G6PC1 structure model identifies regions of the highest predictive confidence (left) and lowest predictive confidence (right) for the “likely benign” and “likely pathogenic” classes. The data identified as highest confidence and lowest confidence were taken from top 10% of scores representing the smallest range and greatest range, respectively, in **(B)**. **F.** Profile of AlphaMissense scores as a function of allele frequency for 493 variants obtained from either the gnomAD_v4.1 or RGC databases. The dashed line indicates the “benign” cutoff for training the AlphaMissense model. Despite genetic evidence linking SNVs at position 295 as causative for GSD1a, AlphaMissense classifies 295 variants as “ambiguous” due to high score variability combined with intermediate levels of sequence conservation **(D)**.

**Figure S2.**
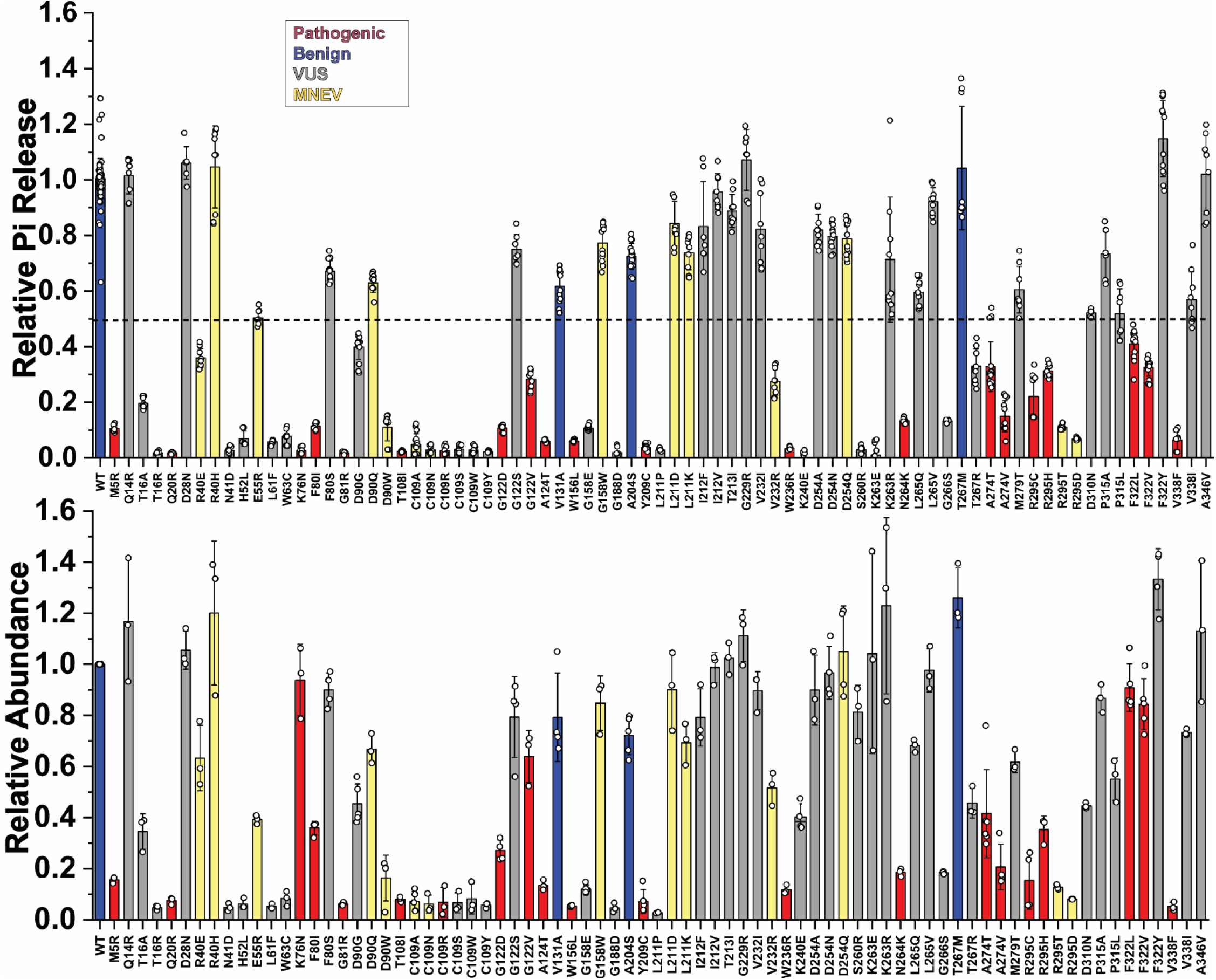
Activity (top) and abundance (bottom) profiles for 78 variants in this study. Both readouts are normalized to the WT. The coloring of each variant is based on the classification scheme as described in Figure 1 of the main manuscript. The horizontal dashed line represents the 50% activity cutoff to preclude the GSD1a phenotype based on the biochemical properties of the variants described here.

**Figure S3.**
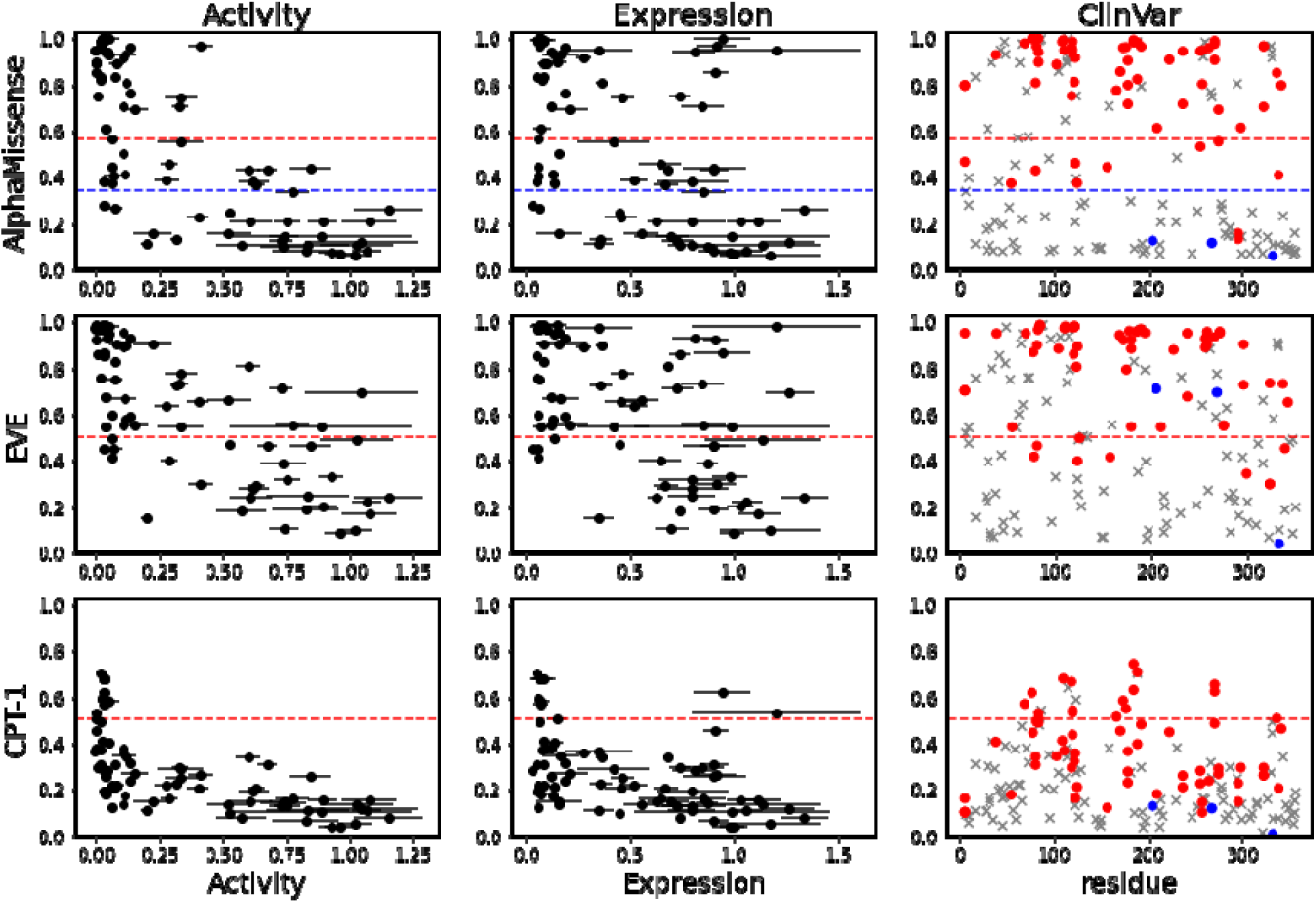
Comparison of variant effect predictors. Despite distinct model designs, similar correlations between AlphaMissense, EVE and CPT-1 scores and biological metrics (Activity and Abundance) were observed. However, AlphaMissense demonstrated greater accuracy in variant classifications of ClinVar variants (far right panels). Pathogenic, red; benign, blue; VUS, grey. Dashed lines represent the classification cutoff.

**Figure S4.**
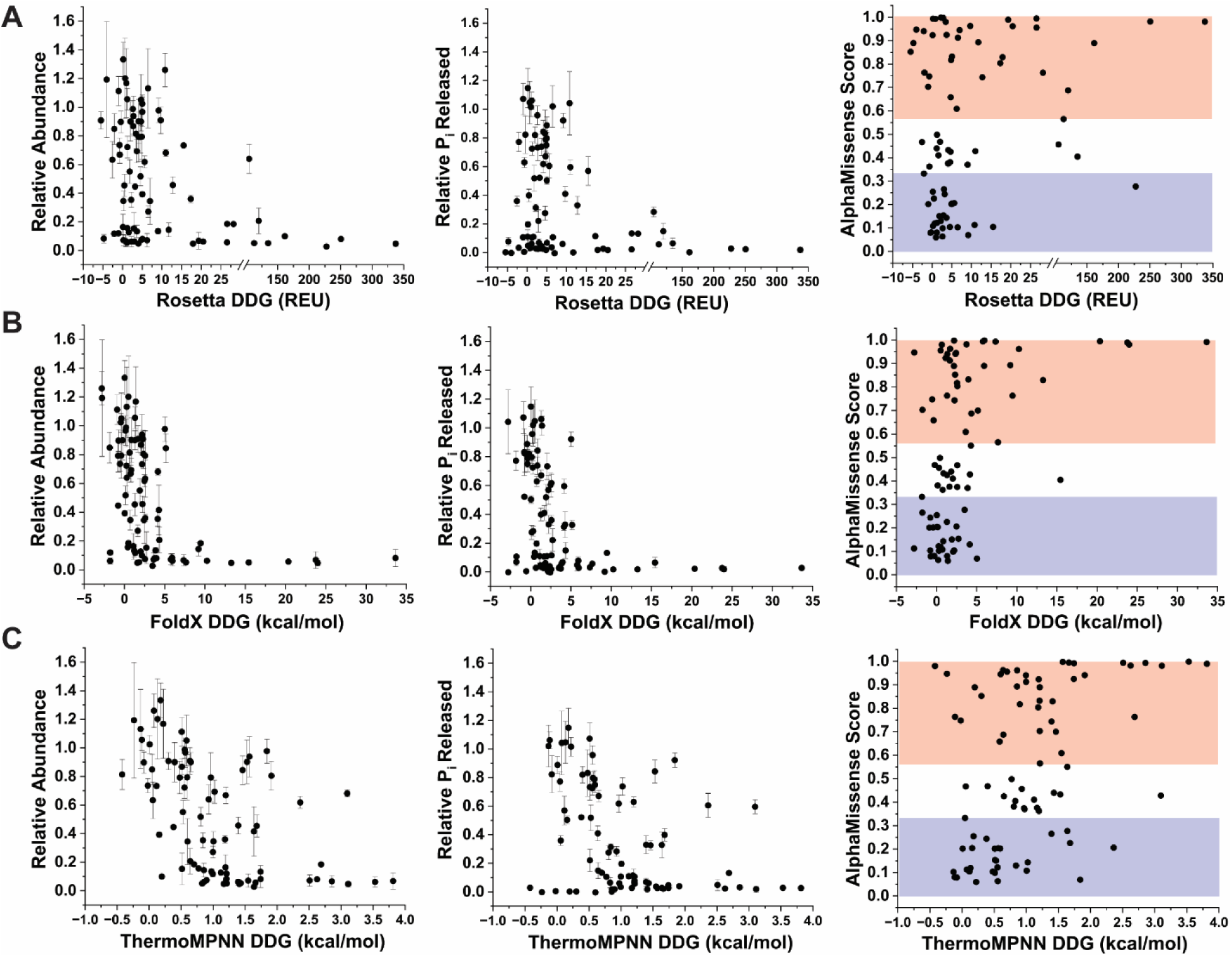
Comparison of ΔΔG stability predictors. Predicted changes in variant stability are shown for Rosetta **(A)**, FoldX **(B)** and the deep learning model ThermoMPNN **(C)** relative to enzyme abundance (left) and activity (center). The right panel for each group shows the relationship between AlphaMissense score and predicted ΔΔG.

**Figure S5.**
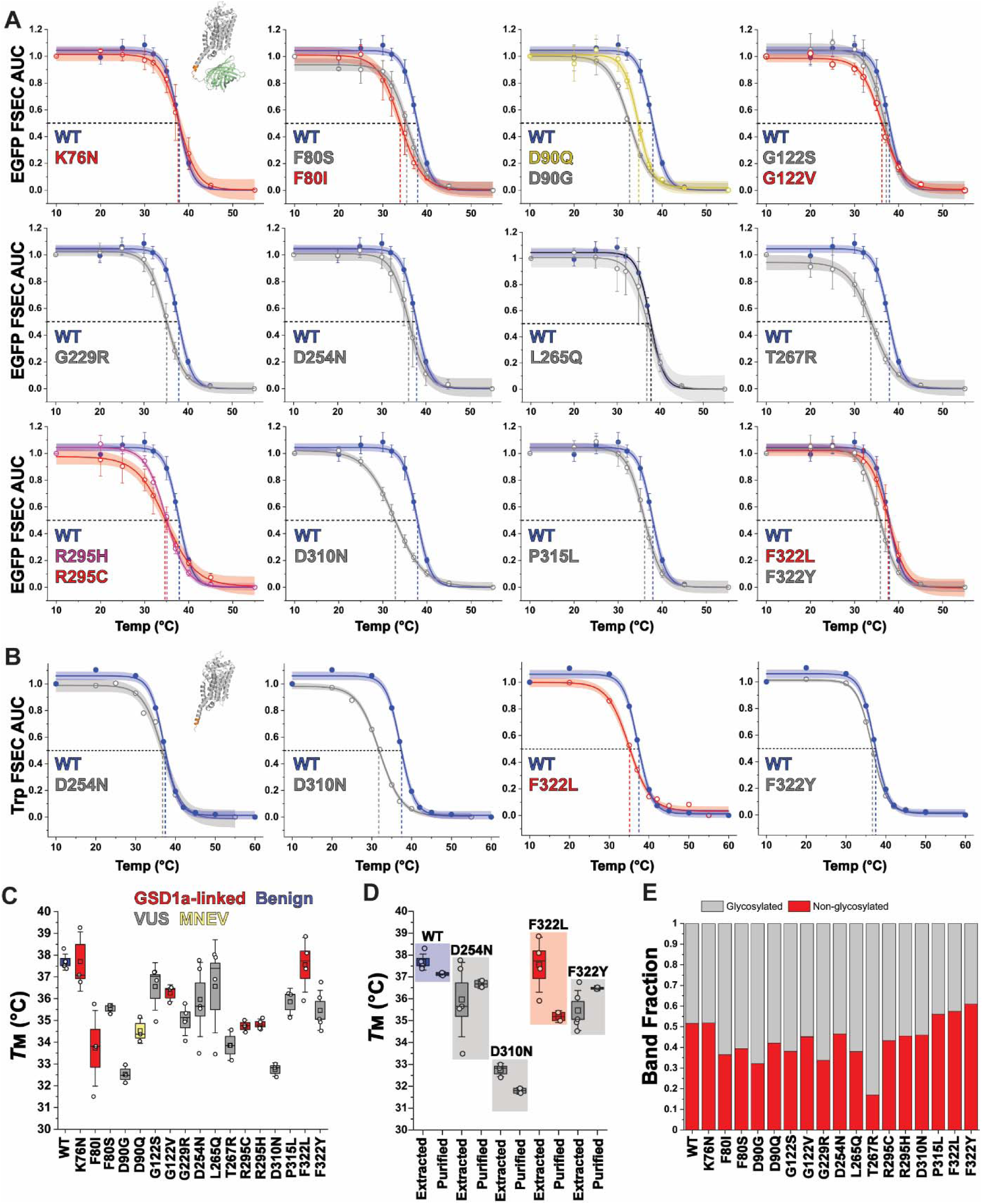
Thermostability analysis of select G6PC1 variants. **A.** Melt curves of non-purified (detergent extracted) G6PC1 obtained from EGFP fluorescence detection size exclusion chromatography illustrate changes in melting temperature indicated by the dashed lines. **B.** Melt curves of purified G6PC1 devoid of the EGFP-His_8_ C-terminal tag. The solid lines through the data points are fits to a dose response model shown with 95% confidence intervals. Melt curves in **(A)** were constructed from n≥3 biological replicates with the mean ± standard deviation. Melt curves in **(B)** were constructed from two biological replicates. **C.** The distribution of *T*_M_ determined from the curves are shown as box plots. **D.** Comparison of *T*_M_ from non-purified and purified samples. The box size in **C** and **D** captures the standard error while the whisker shows the standard deviation. The mean is represented by an open box and the median a solid line. **E.** Relative populations of glycosylated and non-glycosylated G6PC1 determined from western blots using anti-GFP primary antibody.

**Figure S6.**
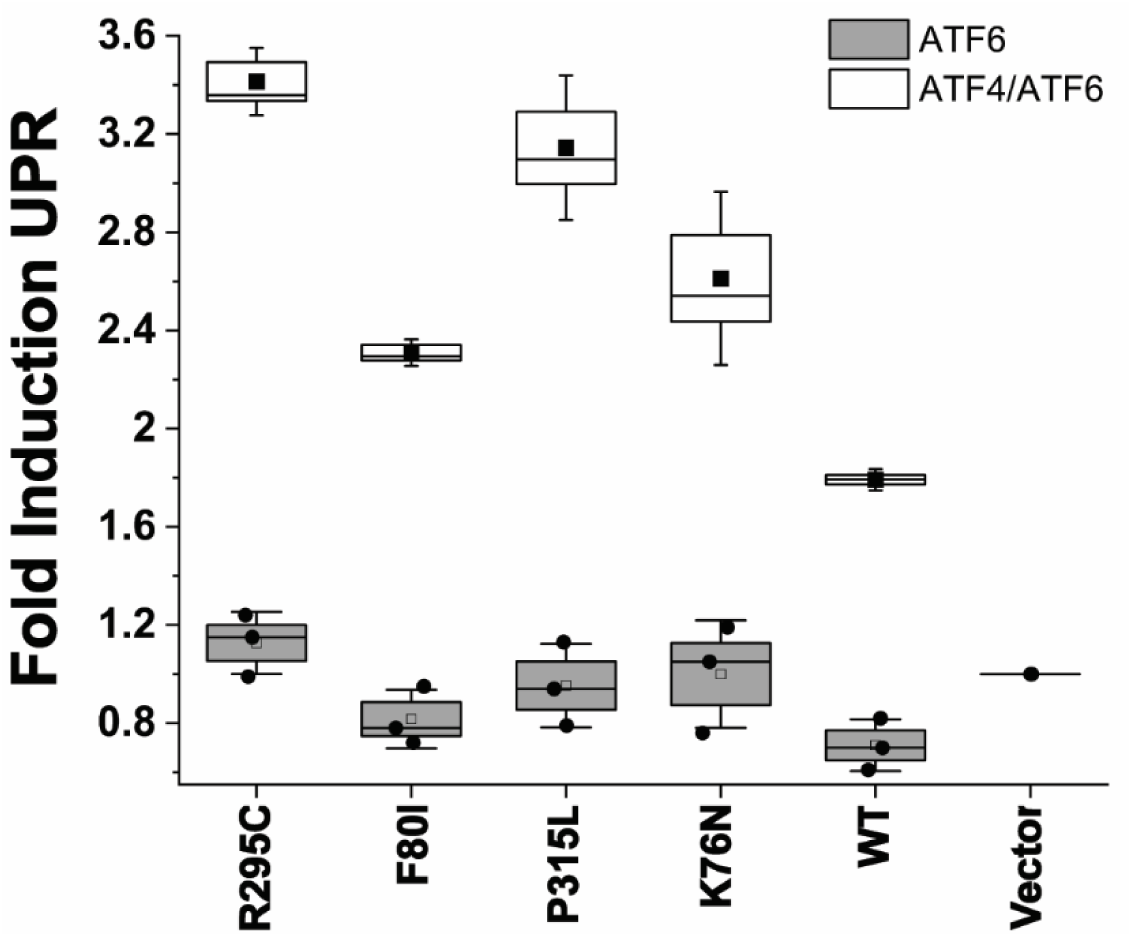
Fine tuning the detection of the unfolded protein response induced by G6PC1 variants. A multimerized ATF4/ATF6 luciferase reporter increased the sensitivity of detection relative to ATF6 alone. The box plots represent the distribution of data collected for three independent experiments. The box size represents the standard error and the whisker the standard deviation. Open/filled squares and the solid line illustrate the mean and median, respectively.

**Figure S7.**
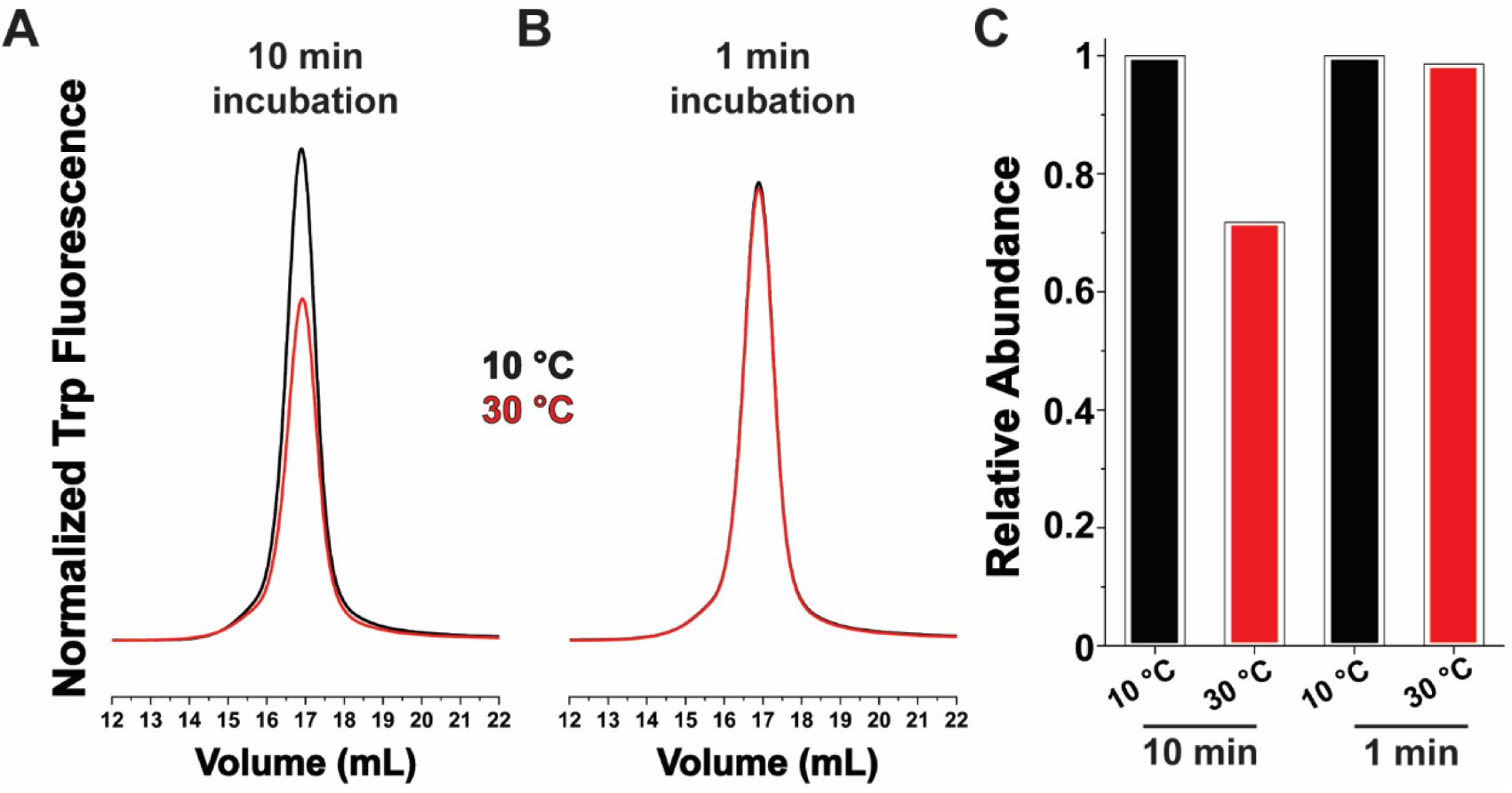
Relative stability of D310N at two incubation temperatures. **A.** Relative to 10 °C, incubation for 10 min at 30 °C induces a loss of folded protein based on Trp fluorescence detection size exclusion chromatography. **B.** There is no observable loss of D310N when incubating at the same temperatures but for only 1 min, which mimics the steady state kinetics assay conditions. **C.** Quantification of the area under the elution curve showing a loss of ∼30% following 30 °C for 10 min.

**Figure S8.**
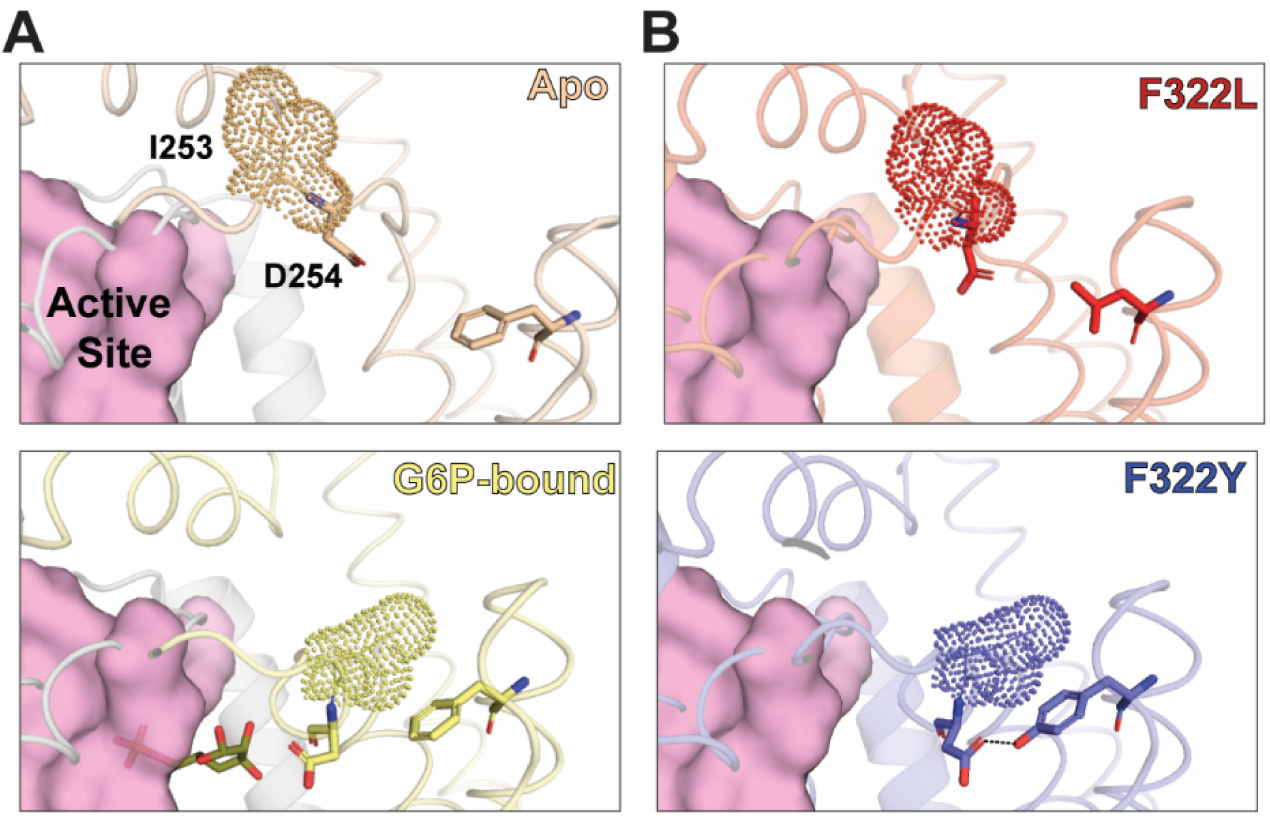
Structural comparison of cryo-EM and AlphaFold models. **A.** The cryo-EM structure of human G6PC1 (PDB ID 9JTL) determined in the absence of substrates (Apo) shows an accessible active site mediated by disengagement of the TM6/7 loop. In contrast, this loop collapses over the active site and anchored by interactions between I253 and F322 in the G6P-bound state (PDB ID 9JTM). **B**. AlphaFold modeling of the F322L variant recapitulates the Apo state cryo-EM structure due to the loss of hydrophobic interactions with I253. In contrast, the F322Y variant stabilizes the closed conformation through a new H-bond between the –OH moiety of Y322 and carboxyl moiety of D254.

